# MAGE-A4-Responsive Plasma Cells Promote Non-Small Cell Lung Cancer

**DOI:** 10.1101/2024.07.10.602985

**Authors:** Dominique Armstrong, Cheng-Yen Chang, Monica J. Hong, Linda Green, William Hudson, Yichao Shen, Li-Zhen Song, Sheetal Jammi, Benjamin Casal, Chad J. Creighton, Alexandre Carisey, Xiang H.-F. Zhang, Neil J. McKenna, Sung Wook Kang, Hyun-Sung Lee, David B. Corry, Farrah Kheradmand

## Abstract

Adaptive immunity is critical to eliminate malignant cells, while multiple tumor-intrinsic factors can alter this protective function. Melanoma antigen-A4 (MAGE-A4), a cancer-testis antigen, is expressed in several solid tumors and correlates with poor survival in non-small cell lung cancer (NSCLC), but its role in altering antitumor immunity remains unclear. We found that expression of MAGE-A4 was highly associated with the loss of *PTEN*, a tumor suppressor, in human NSCLC. Here we show that constitutive expression of human *MAGE-A4* combined with the loss of *Pten* in mouse airway epithelial cells results in metastatic adenocarcinoma enriched in CD138^+^ CXCR4^+^ plasma cells, predominantly expressing IgA. Consistently, human NSCLC expressing MAGE-A4 showed increased CD138^+^ IgA^+^ plasma cell density surrounding tumors. The abrogation of MAGE-A4-responsive plasma cells (MARPs) decreased tumor burden, increased T cell infiltration and activation, and reduced CD163^+^CD206^+^ macrophages in mouse lungs. These findings suggest MAGE-A4 promotes NSCLC tumorigenesis, in part, through the recruitment and retention of IgA^+^ MARPs in the lungs.

## Introduction

Despite the discovery of a growing number of targetable molecular markers associated with non-small cell lung cancer (NSCLC), minimal improvements have been made in the overall survival of patients with advanced disease (1). Small-molecule inhibitors targeting specific mutations have shown only limited short-term clinical efficacy (2, 3). Immunotherapy has further improved outcomes in lung cancer patients, but the objective response rate in non-selected patients has remained low (4). Checkpoint blockade has illuminated the interplay between malignant cells, their microenvironment, and the critical importance of the adaptive immune cells infiltrating tumors. Furthermore, several animal models of NSCLC using mutations commonly seen in patients (e.g., *Kras, Egfr, Pten, Smad4,* and *Lkb1*) have provided important advances in deciphering how individual mutations affect the tumor immune microenvironment (TIME) (5–12). However, the incomplete response rates of current therapies reveal an unmet need for a more comprehensive understanding of how additional tumor-specific factors alter their locoregional immune microenvironment (5).

Tumor-intrinsic factors such as cancer-testis antigens (CTAs) are expressed in several solid tumors and have been evaluated as therapeutic targets or used for prognostication (13–19). However, little is known about CTAs’ roles in promoting tumors and/or altering the TIME. CTAs are restricted to the testis and placenta, suggesting they have a functional role in cellular physiology, while their restrictive expression among different stages of development and germ cells indicates a high degree of regulation (20). Melanoma antigen A4 (MAGE-A4), a prototypic CTA, is commonly expressed in NSCLC, and negatively correlates with 5-year overall survival even in those with early-stage disease (18), indicating its role in promoting tumor growth and invasion. MAGE proteins are considered immunogenic, as antigen-specific humoral and cellular immunity have been detected within tumors and in the peripheral circulation (21–25). However, whether and how MAGE-A4 can promote NSCLC or alter the TIME has not been examined.

The implementation of immune checkpoint inhibitors (ICIs), a paradigm shift in cancer treatment, has provided a stable response for a subset of patients with NSCLC (4). However, tumor-specific immune-modulating factors could subvert antitumor immunity beyond the targeted T cell signature checkpoint ligands underlining the need to investigate mechanisms of treatment resistance. Specifically, poorly immunogenic (e.g., cold) tumors show reduced T cell infiltration and low program-death ligand-1 (PD-L1) expression, while Tregs, myeloid-derived suppressor cells (MDSCs), and tumor-associated macrophages can infiltrate and suppress cytolytic CD8^+^ T cell responses (5, 26). In contrast, highly immunogenic (e.g., hot) tumors are characterized by an abundance of T lymphocytes (5), along with other immune cells, which vary by tumor type. This complex milieu raises the question of whether the intrinsic properties of neoplastic cells that generate CTAs could govern the immune cells within the TIME.

Most studies of tumor-related immune responses are focused on T lymphocytes because B cells are not the most abundant leukocytes within the lung TIME. However, B cells represent some of the most altered immune cells in human NSCLC (27, 28). The abundance of B cells in tertiary lymphoid structures correlates with improved survival (21); however, a role for the immunosuppressive subset of B cells, B-regulatory cells, has also been described in NSCLC and other solid tumors (29, 30). Similarly, dual roles in tumor development or antitumor immunity for plasma cells (terminally differentiated B cells) have been described (30–32). Specifically, plasma cells have been described as a dominant infiltrating cell in some NSCLC (33, 34), and have shown both pro- and antitumor associations in several clinical studies (33, 35–39). One retrospective study suggested that an increased abundance of plasma cells in the absence of other B cell subsets is associated with poor outcomes (40). These findings indicate that tumor intrinsic factors may be critical determinants of whether plasma cells are beneficial or detrimental in lung cancer development.

Here, we investigated the malignant potential and immunological roles of MAGE-A4 in mouse and human NSCLC. Using The Cancer Genome Atlas (TCGA), we identified a correlation between MAGE-A4 expression and multiple aberrations in the tumor suppressor, *PTEN*. We then developed an autochthonous mouse model in which human MAGE-A4 is constitutively expressed in mouse lung epithelial Club cells. Airway-specific expression of MAGE-A4 along with deletion of *Pten* resulted in invasive lung adenocarcinoma that spontaneously metastasized to distant organs. We found an accumulation of CD138^+^ IgA^+^ plasma cells in mouse and human tumors that expressed MAGE-A4, which we termed <u>M</u>AGE-<u>A</u>4-responsive <u>p</u>lasma cell<u>s</u> (MARPs). Here, we demonstrate that MAGE-A4 is an oncoprotein that potentiates NSCLC progression by recruiting pro-tumorigenic plasma cells in the tumor microenvironment to drive tumor invasion and metastasis.

## Results

### MAGE-A4 is expressed in human squamous and adeno subtypes of NSCLC

MAGE-A4 is highly prevalent in NSCLC (16–19). Interrogating TCGA, we found that MAGE-A4 expression showed large variation depending on the stage in NSCLC (n=989) (Figure 1A). To further characterize MAGE-A4, we performed immunohistochemistry (IHC) using an array of randomly selected tissue from 29 different cases of NSCLC that included the subtypes lung adenocarcinoma (LUAD), lung squamous cell carcinoma (LUSC), mixed features (e.g., adenosquamous) and large cell; these latter 3 types we grouped as LUSC/mixed (Supplemental Table 1). In one case, the selected region to generate the tissue array lacked tumor cells, thus it was excluded from our analysis. Ten tumors (∼36%) were MAGE-A4 positive (MAGE-A4^pos^); specifically, 5/11 (∼45%) LUAD and 5/17 LUSC/Mixed (∼29%) expressed MAGE-A4 (Figure 1B and 1C). All 10 MAGE-A4^pos^ tumor samples showed nuclear staining, while 4 had additional cytoplasmic expression; 1/5 LUAD exhibited nuclear with cytoplasmic MAGE-A4 expression while 3/5 in LUSC/Mixed show this pattern (Figure 1C). Given the heterogeneity of MAGE-A4 in NSCLC, we next questioned whether its expression could be associated with alterations in specific tumor suppressor genes. We found a significant correlation between MAGE-A4 expression with reduced copy number, deletion, and somatic mutation of *PTEN* mRNA in several LUSC (SQ.1, SQ.2a, SQ.2b) and in LUAD (AD.1) molecular subtypes in TCGA database (41) (Figure 1D). Using genomics data from an independent cohort, the Clinical Proteomic Tumor Analysis Consortium dataset, we validated the significant correlation between *MAGE-A4* expression and *PTEN* loss in NSCLC (Supplemental Figure 1). These findings suggest that MAGE-A4 is prevalent in many types of human NSCLC and its expression is associated with PTEN aberrations.

**Fig. 1:**
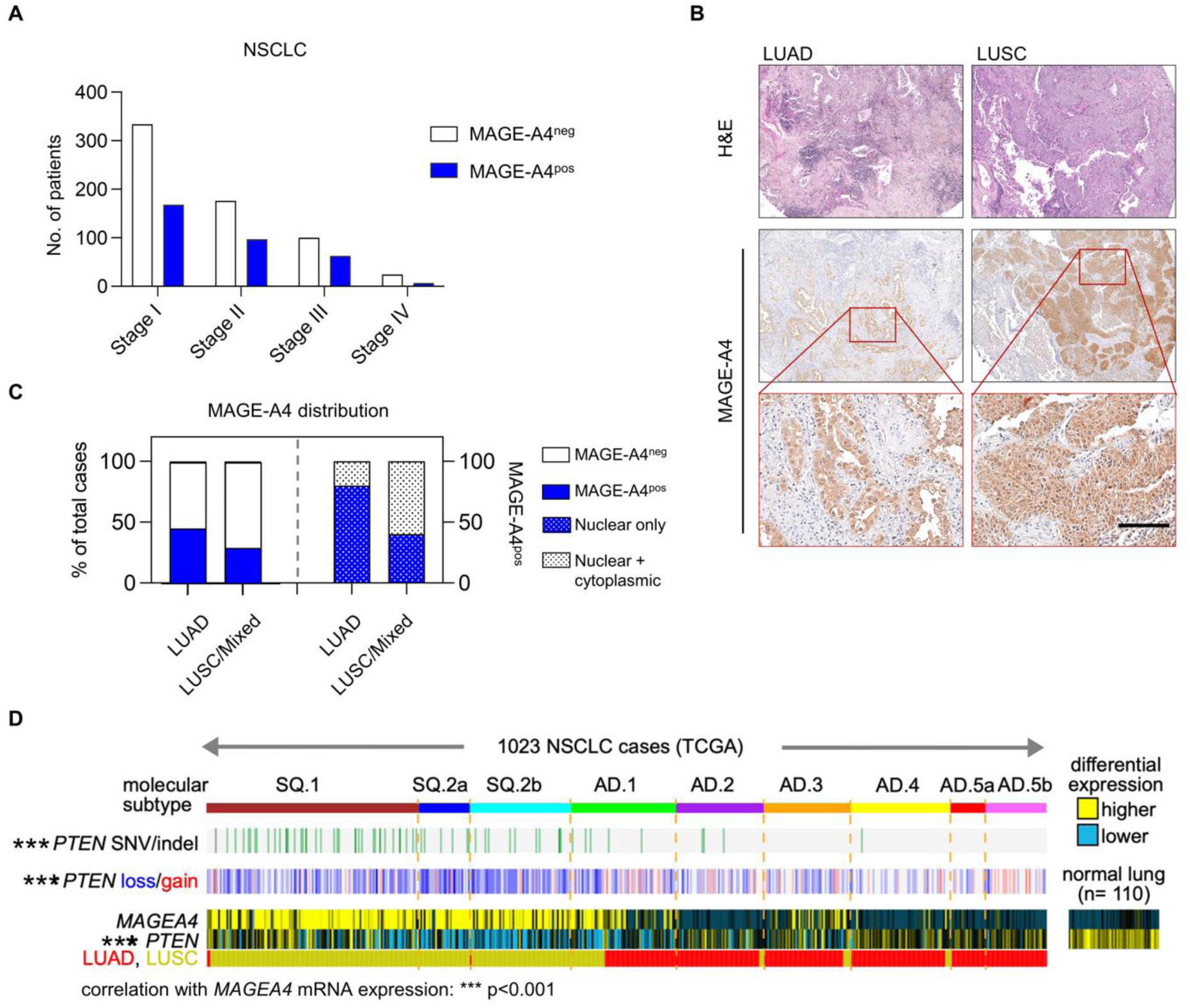
MAGE-A4 is expressed in human NSCLC and is associated with loss of *PTEN*. (**A**) MAGE-A4 expression in NSCLC by RNA from TCGA dataset, n=989. FPKM ≥ 1 was considered positive. (**B**) Representative H&E of human NSCLC and serial sections show MAGE-A4^pos^ staining (IHC) in lung adenocarcinoma (LUAD) and in lung squamous cell carcinoma (LUSC); 100x (top), scale bar 150 μm (insets) (**C**) Incidence of MAGE-A4 in NSCLC array of 28 NSCLC tumors and cellular localization of MAGE-A4 as determined by IHC detection. LUSC/Mixed includes LUSC, adenosquamous (mixed features), and large-cell carcinoma subtypes. Total MAGE-A4^pos^ in tumor cases (left graph) and all MAGE-A4^pos^ by subcellular localization (right side of graph) (**D**) The Cancer Genome Atlas (TCGA) data of 1023 cases shows MAGE-A4 expression in human NSCLC is associated with decreased *PTEN*. Association of MAGEA4 with *PTEN* mRNA and copy levels by Pearson’s correlation (*P*=0.00098 and *P*=1.3E-7, respectively); association of MAGEA4 with *PTEN* mutation, by t-test (*P*=1.4E-6)

### Airway-specific expression of human *MAGE-A4* combined with *Pten* loss causes invasive lung adenocarcinoma

Because MAGE-A4 expression in multiple subtypes of NSCLC was significantly associated with the loss of PTEN, we next sought to determine its role in lung cancer development and progression. To address this question, we engineered a human (h)*MAGE-A4* construct under the LoxP-Stop-LoxP to generate its conditional expression in mice (*MAGE-A4^LSL^*) (Supplemental Figure 2, A-C). We crossed *MAGE-A4^LSL^* mice to Club cell-specific promoter iCre mice (*CCSP^iCre^*) (42) to generate a new model of bigenic *CCSP^iCre^MAGE-A4^LSL^* (*MAG4*) mice. Notably, *MAG4* mice were born with normal genetic frequency and did not develop tumors (Supplemental Figure 2D). To examine the significance of MAGE-A4 expression with *PTEN* loss, as we discovered in our in silico human data analyses, we crossed *MAG4* to *Pten*-floxed mice (42) to generate *CCSP^iCre^ MAGEA4^LSL^ Pten^f/f^* (*MAG4/Pt*) mice (Figure 2A). We found that much like *Cre^-^MAG4/Pt* (*CCSP^WT^ MAGEA4^LSL^ Pten^f/f^*) used as wild-type controls, neither h*MAGE-A4* expression nor deletion of *Pten* alone in Club cells (*CCSP^iCre^ Pten^f/f^*; *Pt*, Figure 2A) was sufficient to drive lung tumors (Figure 2B, Supplemental Figure 2D). While some studies have shown that at 12 months, 25% of mice with *Pten* deletion in airways develop non-invasive lung cancer (43), *MAG4/Pt* mice developed invasive tumors in the lung as early as 3 months of age, with 100% developing tumors by 5 months of age (Figure 2B and C, Supplemental Figure 2E). The cells showed invasive features as their basement membrane was disrupted, and morphologically cells showed distinct high nuclear-to-cytoplasmic content, were heterogeneous, and disorganized compared with neighboring cells (Figure 2, B and D). We confirmed that hMAGE-A4 is not expressed in *Cre^-^MAG4/Pt* or *Pt* lung, but *MAG4* and *MAG4/Pt* mice exclusively express cytoplasmic and nuclear hMAGE-A4 in the bronchial epithelial cells (Figure 2D), as we had detected in human tumors (Figure 1C). *MAG4/Pt* tumors initiate in the airway and resemble an acinar adenocarcinoma subtype as they formed ductal-like patterns and expressed TTF1 and KRT7 while lacking squamous carcinoma markers KRT5 and SOX2 (42) (Figure 2, B and D, Supplemental Figure 3). Notably, *MAG4/Pt* tumors spontaneously metastasized to the spleen, intestines, and cervical lymph nodes at 5, 10, and 13 months of age, respectively (Figure 2, E and F, Supplemental Figure 4, A-D). *MAG4/Pt* mice develop enlarged spleens, with disrupted architecture including displaced germinal centers, and are positive for MAGE-A4, compared with *Cre-MAG4*/*Pt* mice (Supplemental Figure 4, B-D). We further validated tumor micro-metastasis by detecting the mRNA expression of *Scgb1a1*, a Club-cell-specific gene, in the spleen (Figure 2G). Together, these results demonstrate that h*MAGE-A4* in the absence of *Pten* promotes NSCLC tumorigenesis and metastasis.

**Fig. 2:**
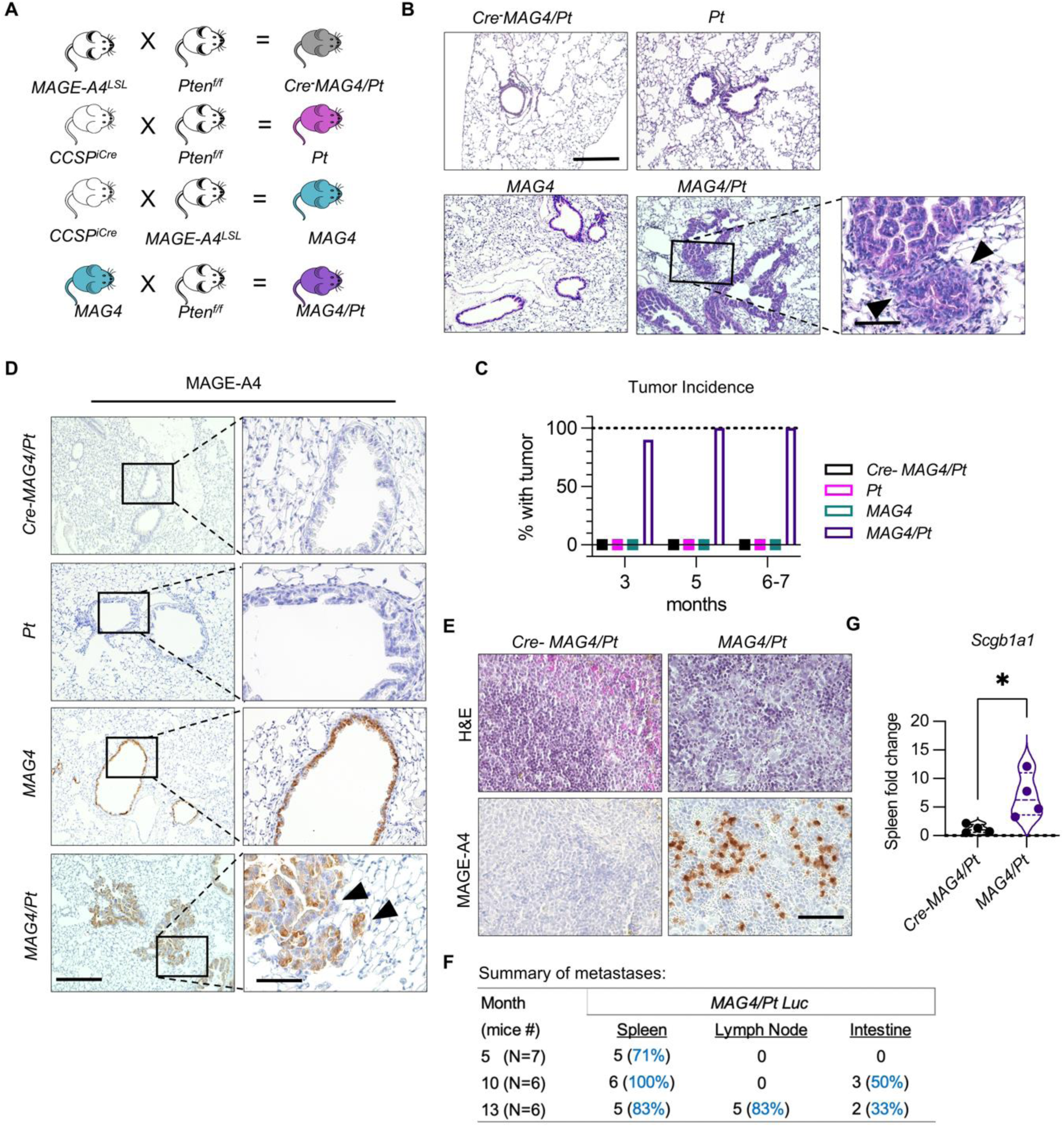
De-novo hMAGE-A4 expression in the lung causes invasive LUAD in the absence of *Pten*. (**A**) Breeding scheme to generate mouse models and controls: *CCSP^WT^ hMAGE-A4^LSL^/Pten^f/f^* (*Cre-MAG4/Pt*), *CCSP^iCre^ Pten^f/f^* (*Pt*), *CCSP^iCre^ hMAGE-A4^LSL^* (*MAG4*), and *CCSP^iCre^ hMAGE-A4^LSL^*/*Pten^f/f^* (*MAG4/Pt*). (**B**) Representative H&E-stained images of *Cre-MAG4/Pt, Pt*, *MAG4*, and *MAG4/Pt* lung from 6-7-month-old mice demonstrate tumors in *MAG4/Pt* but not in *MAG4* or *Pt* alone. Arrows indicate areas with loss of basement membrane and invasion. The scale bar is 300 μm; the inset scale bar is 150 μm. (**C**) Tumor incidence among groups of mice at indicated ages as determined by histology. n=8-27 per group. (**D**) Representative IHC of MAGE-A4 expression in all four genotypes at 6-7 months of age. Black arrows in *MAG4/Pt* indicate areas of invasion with loss of basement membrane. Scale bar is 300 μm (left panel), 75 μm (right insets) (**E**) H&E (top) and respective IHC of MAGE-A4 (bottom) of *Cre-MAG4/Pt* and *MAG4/Pt* spleen at 12 months of age, scale bar 75 μm. (**F**) Summary of *MAG4/Pt* tumors metastasize to spleen, intestine, and cervical lymph nodes by age as determined from D-luciferin screening. (**G**) Detection of *Scgb1a1* in 12-13 months in *Cre-MAG4/Pt* vs. *MAG4/Pt* spleen by qPCR, *n* = 4 per group, one experiment. Student’s t-test, \**P* <0.05

### Increased CD163^+^ CD206^+^ macrophages and reduced CD103^+^ APCs in *MAG4/Pt* mice

The TIME can play opposing roles by either promoting or eliminating malignant cells (5). Therefore, we next examined the innate immune cells in the lungs of *MAG4/Pt* mice at 5 months of age, when 100% develop tumors with ∼70% showing metastasis (Figure 2C and F). We found that lung F4/80^+^ CD64^+^ macrophages were increased in *Pt* mice but were significantly reduced in *MAG4/Pt* mice (Supplemental Figure 5, A and B). However, *MAG4/Pt* mice showed an expansion of tumor-associated macrophages expressing CD163^+^ CD206^+^ in the lungs (Supplemental Figure 5C). We next examined CD103^+^ antigen-presenting cells, which are critical in cross-presenting tumor antigens to CD8 T cells and play a key role in antitumor immunity (44). We found that CD103^+^ DCs were increased in the lungs of *Pt* mice, but were significantly reduced in *MAG4/Pt* mice, demonstrating that the expansion of this antitumor cell population is driven by loss of airway *Pten* (Supplemental Figure 5D). Neither Ly6C^+^ mononuclear (M) myeloid-derived suppressor cells (M-MDSCs) nor Ly6G^+^ polymorphonuclear (PMN-MDSCs), were altered in *MAG4/Pt* (Supplemental Figure 5, E-H). These findings suggest that hMAGE-A4 alters lung myeloid populations by reducing antigen-presenting cells while promoting an increase in tumor-associated macrophages in the lungs.

### Reduced CD4^+^ T cells and lack of T cell infiltration in *MAG4/Pt* lungs

We next interrogated the adaptive immune microenvironment in *MG4/Pt* mice at 5 months of age. We found a decrease in the total number of CD4^+^ T cells and CD25^+^ FOXP3^-^ activated subset in *MAG4/Pt* mice compared with *Pt* (Supplemental Figure 6, A-D). Unexpectedly, total T cells and other CD4^+^ T cell subsets including Tregs (FOXP3^+^), activated Tregs (CD25^+^ FOXP3^+^), T helper type1 (Th1, T-bet^+^) and T helper type 17 (Th17, RORγT^+^) were not increased in the lungs of *MAG4* or *MAG4/Pt* (Supplemental Figure 6B and E-H). Furthermore, CD8 T cells and activated subsets (CD25^+^, T-bet^+^) (Supplemental Figure 6I to K) were also not significantly altered in mice expressing hMAGE-A4. While we found a lack of T cell activation, there was an increase in T helper Type 2 (Th2, GATA3^+^) that approached significance (Supplemental Figure 6L). Remarkably, exhausted PD-1^+^ CD4 or PD-1^+^ CD8 T cells were unaltered in *MAG4/Pt* lung compared with *Cre^-^MAG4/Pt* (Supplemental Figure 7A, C, and I). Furthermore, CD4 or CD8 T cells expression levels of PD-1 co-inhibitory molecules (CD38, LAG3, TIM3, CTLA4) were unchanged in *MAG4/Pt* lung (Supplemental Figure 7, B-M). Collectively, these findings suggest a role for hMAGE-A4 in reducing antitumor immunity through a reduction in the total number of CD4^+^ T cells and the CD25^+^ FOXP3^-^ activated subset.

### Plasma cells accumulate in *MAG4/Pt* lung

Histological evaluation of *MAG4/Pt* mice lungs consistently showed dense infiltration of immune cells with eosin-stained cytoplasm (indicative of protein synthesis), distinguishing perinuclear hof, and clockface, eccentric nuclei (45) (Figure 3A). Because these morphologic characteristics were suggestive of plasma cells, we confirmed their identity using CD138, a well-defined plasma cell marker (46), and CXCR4, the key chemokine receptor known to direct migration of plasma cells (47) (Figure 3, B and C, Supplemental Figure 8, A-C). Consistent with CD138^+^ and CXCR4^+^ IHC staining, flow cytometry confirmed CD138^+^ CXCR4^+^ plasma cells were increased in *MAG4/Pt* lung (Figure 3D). Because CD138^+^ CXCR4^+^ plasma cells were only observed in lung tumors expressing MAGE-A4, we termed them <u>M</u>AGE-<u>A</u>4-responsive <u>p</u>lasma cell<u>s</u> (MARPs). Further analysis of MARPs confirmed the expression of IgA, IL-10, and TGF-β (Figure 3, D-G, Supplemental Figure 9A), which are associated with an immunosuppressive phenotype (32, 48). Consistently, IgA antibody isotype was significantly increased in bronchial alveolar lavage (BAL) fluid, while other isotypes were unaltered (Supplemental Figure 9, B-G). Since IgA isotype requires class-switching, which involves T follicular-helper (Tfh) cells in mature germinal centers (49, 50), we evaluated Tfh cells in *MAG4/Pt* mice. Tfh cells were not significantly increased (Supplemental Figure 9H), indicating that in this model, MARPs are expanded in a T-cell-independent manner (51).

**Fig. 3:**
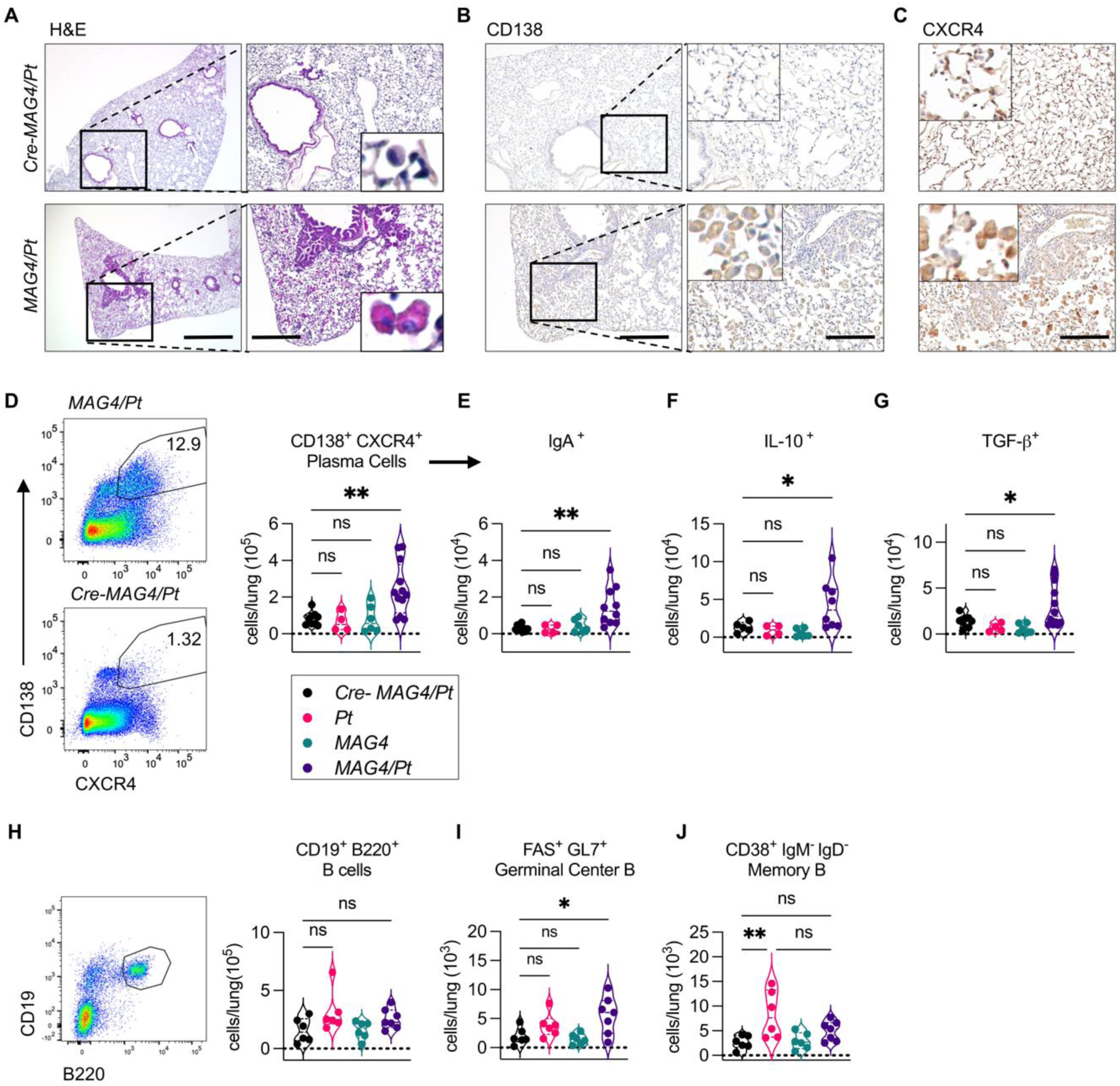
*MAG4/Pt* mouse lungs accumulate plasma cells. (**A**-**C**) Representative serial section lung 5-month-old *MAG4/Pt* and *Cre-MAG4/Pt* mouse. (**A**) H&E, scale bar 770 μm (left) and 300 μm (right) (**B**) IHC of CD138 serial section from (A), scale bars 300 μm (left) and 150 μm (right). (**C**) IHC of CXCR4 serial section from (A), scale bar: 150 μm. (**D**) Gating and quantification of plasma cells (EpCAM^-^ CD138^+^ CXCR4^+^) from 5-6-month-old mouse lung by flow cytometry. (**D**-**G**) Quantification of (**E**) IgA, (**F**) IL-10, and (**G**) TGF-β-expressing plasma cells in cells from panel (D) Two experiments combined, n=4-10 per group (**H**-**J**) Flow cytometry of B cell subsets (**H**) Gating and quantification of CD19^+^ B220^+^ B cells (**I**,**J**) Quantification of CD19^+^ B220^+^ subsets (**I**) FAS^+^ GL7^+^ germinal center B cells and (**J**) CD38^+^ IgM^-^ IgD^-^ Memory B cells. One experiment, n= 5-7 per group. Significance determined by One-way ANOVA with Tukey’s or Dunnett’s correction. ns= not significant, \**P* < 0.05, \*\**P* < 0.01.

Given the increase in MARPs, we assessed whether other B cell populations were altered in *MAG4/Pt* lung tumors. While total B220^+^ B cells were not increased in any of the groups, germinal center B cells (GL7^+^ FAS^+^ CD19^+^ B220^+^) were increased in *MAG4/Pt* lung (Figure 3, H and I, Supplemental Figure 9I), suggesting activation of antigen-specific responses. Notably, *Pt* mice showed an increase in memory B cells (CD38^+^ IgD^-^ IgM^-^ CD19^+^ B220^+^) which were suppressed in *MAG4/Pt* (Figure 3J, Supplemental Figure 9I). Cumulatively, these results show that MARPs are the predominant B cell subset most altered in response to hMAGE-A4 expression.

### Single-cell RNA seq substantiates altered B cell subsets in *MAG4/Pt* lungs

To further interrogate the impact of hMAGE-A4 on B cells and plasma cells, we next performed single cell RNA sequencing (scRNA-seq) of whole lung using *Cre-MAG4/Pt*, *Pt*, *MAG4*, and *MAG4/Pt* mice at 5 months of age. We identified epithelial, endothelial, and immune cell clusters using singleR and validated with cell-specific markers (Figure 4A, Supplemental Figures 10 and 11) (52). Consistent with flow data, scRNA-seq showed that overall T cells were not changed, while there were fewer CD8 T cells in *MAG4/Pt* lung (Supplemental Figure 12, A and B). Macrophage subsets were the most altered cell types in *Pt* mice (Supplemental Figure 12C). We next re-clustered B cells and found six distinct B cell subsets: Naïve 1-3, Lambda, Memory, and Plasma cells (PC) (Figure 4B). Consistent with the flow cytometry data, we found B cell clusters in mice expressing hMAGE-A4 were altered because distinct clustering separated *MAG4* and *MAG4/Pt* from the other groups (Figure 4C). Among markers of B cell subsets, *Ighd* and *Ighm* co-expression indicated that they were predominantly naïve B cells (Supplemental Figure 13A). We next identified plasma cells (expressing *Cd9*, *Ly6a*, and *Cd93* and lack of *Pax5*) (53–56), IgM memory B cells (expressing *Ighm*, *Spib*, and lack of *Ighd*), and lambda-light-chain B cell clusters (expressing *Iglc1*) (Figure 4D, Supplemental Figure 13, A and B). Notably, MARPs were exclusively present in the *MAG4/Pt* lungs (Figure 4, B-E). *MAG4* and *MAG4/Pt* mice also showed similar differentially expressed genes (DEGs) in total B cells, indicating that they were transcriptionally distinct from the other two groups (Figure 4F). The highest DEGs in B cells and plasma cells encoded heat-shock proteins, such as Hsp40 and Hsp90 in *MAG4* and *MAG4/Pt* mice. As heat-shock protein genes and *Cd83* were increased and are associated with IL-10 production (57, 58), we evaluated upstream genes that are known to control *Il10* expression. We found increases in several related genes, including *Tgfb1* in B cells in *MAG4* and *MAG4/Pt* (Supplemental Figure 13C). We next confirmed that IL-10 protein is increased in whole-lung homogenate in *MAG4* and *MAG4/Pt* (Supplemental Figure 13D). Together, these findings indicate that hMAGE-A4 is associated with a regulatory-transcriptomic repertoire in B cells in the lung and that plasma cells are uniquely enriched in *MAG4/Pt* mice.

**Fig. 4:**
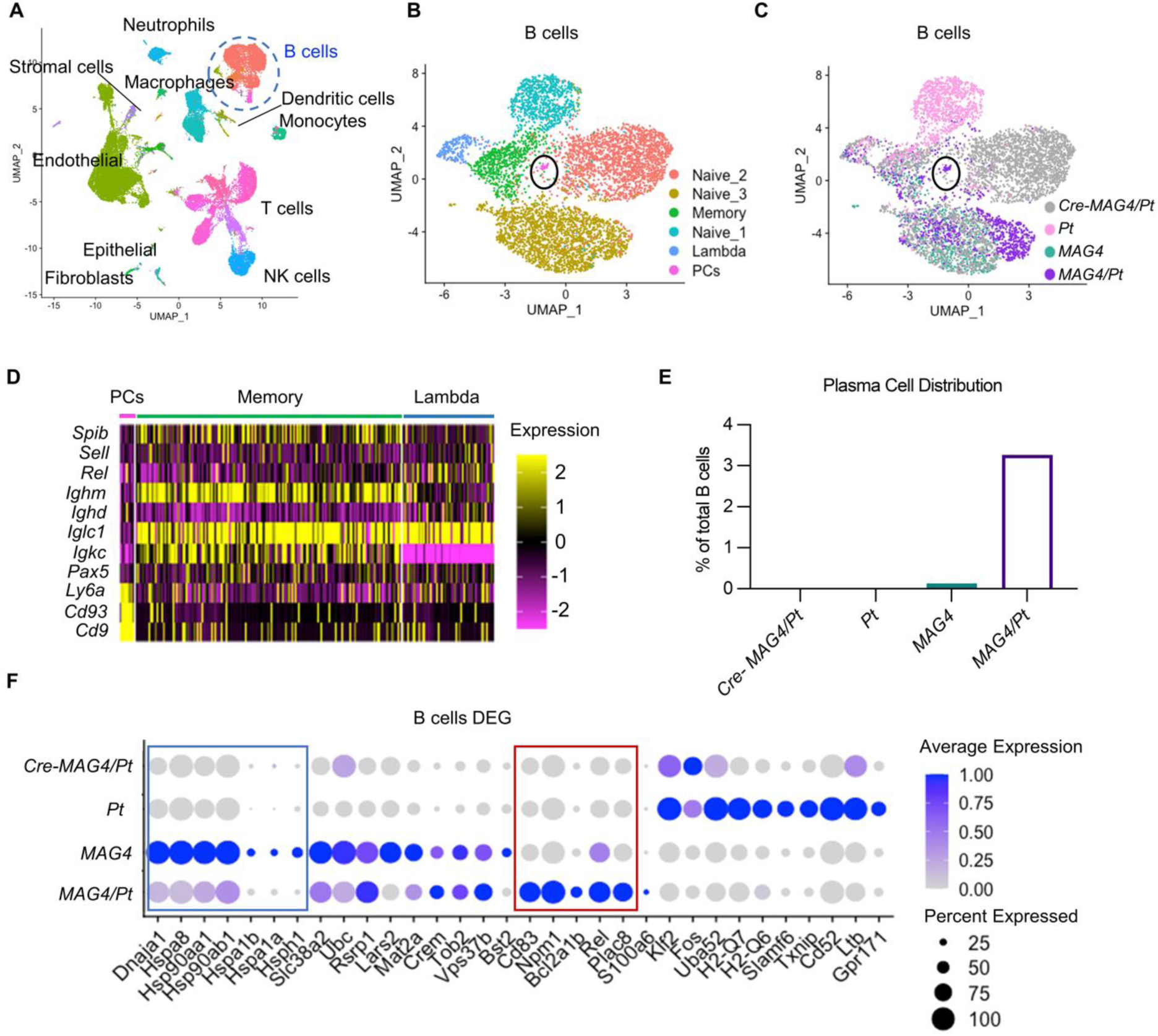
Altered B cell subsets in response to hMAGE-A4 expression in the lungs. (**A**) scRNA-seq of the whole lung identifies multiple cell types by UMAP all 4 genotypes combined (**B**) UMAP of re-clustered B cells from (A) by B cell subsets. (**C**) UMAP of the distribution of B cell subsets by genotypes (**D**) Heatmap of gene expression in distinct B cell subsets: plasma cells (PCs), memory B cells, and λ-light-chain (Lambda) B cells; (**E**) Distribution of plasma cells among groups of mice as a percentage of total B cells (**F**) Dotplot of top differentially expressed genes (DEG) in B cell clusters. The blue box separates heat shock and related proteins; the red box demarcates genes related to the inhibition of apoptosis, autophagy, proliferation, and activation.

### MAGE-A4 promotes plasma cell accumulation via CXCL12/CXCR4 axis

Since MARPs in *MAG4/Pt* mice expressed CXCR4 (Figure 3 A-D), we next measured CXCL12, the key chemokine that recruits plasma cells after differentiation in secondary lymphoid organs (59). As expected, we found that CXCL12 protein was significantly increased in *MAG4/Pt* whole-lung homogenate (Figure 5A). In contrast, CXCL13, a chemotactic factor for B cells (60), was not increased (Supplemental Figure 14A), consistent with our findings that B220^+^ B cells were not increased in *MAG4/Pt* lungs (Figure 3H). We found that *Cxcl12* expression was localized to the lung endothelial cells as evidenced by its overlap with the expression of endothelial cell marker *Pecam1* in scRNA-seq data (Figure 5B). Moreover, an increase in *Cxcl12* in endothelial cells was specific to *MAG4* and *MAG4/Pt* mice (Figure 5C). To functionally validate this observation, we co-cultured human endothelial cells (HUVECs) with H1299 cells and found a 1.8-fold increase in *CXCL12* when tumor cells expressed MAGE-A4 (Figure 5, D and E). *CXCL12* was undetectable in H1299 cells (Supplemental Figure 14B), confirming that endothelial cells were the source of *CXCL12* expression. Recombinant MAGE-A4 did not increase *CXCL12* in HUVEC cells (Supplemental Figure 14C), indicating the indirect effect of MAGE-A4^pos^ tumors on endothelial cells. To assess a potential mechanism for this interaction, we compared DEGs from *MAG4/Pt* vs. *Cre-MAG4/Pt* endothelial cells in NCI Cancer Pathways (61) and found HIF1⍺ as the top differentially expressed pathway (Supplemental Figure 14D). Because HIF1⍺ can directly bind *CXCL12* promoter (62, 63), we examined whether MAGE-A4^pos^ tumor cells induce *CXCL12* through HIF1⍺. As expected, MAGE-A4^pos^, but not MAGE-A4^k/o^ H1299 cells induced the expression of *CXCL12* in human endothelial cells, which was abrogated by PX-478, a specific HIF1⍺ inhibitor (Supplemental Figure 14E). These findings suggest that MAGE-A4-induced *CXCL12* expression in this system is indirect and requires HIF1⍺. Collectively, these data suggest that MAGE-A4 expression in tumors can augment the production of CXCL12 in endothelial cells and is a plausible mechanism for the retention of MARPs in MAGE-A4^pos^ tumors.

**Figure 5:**
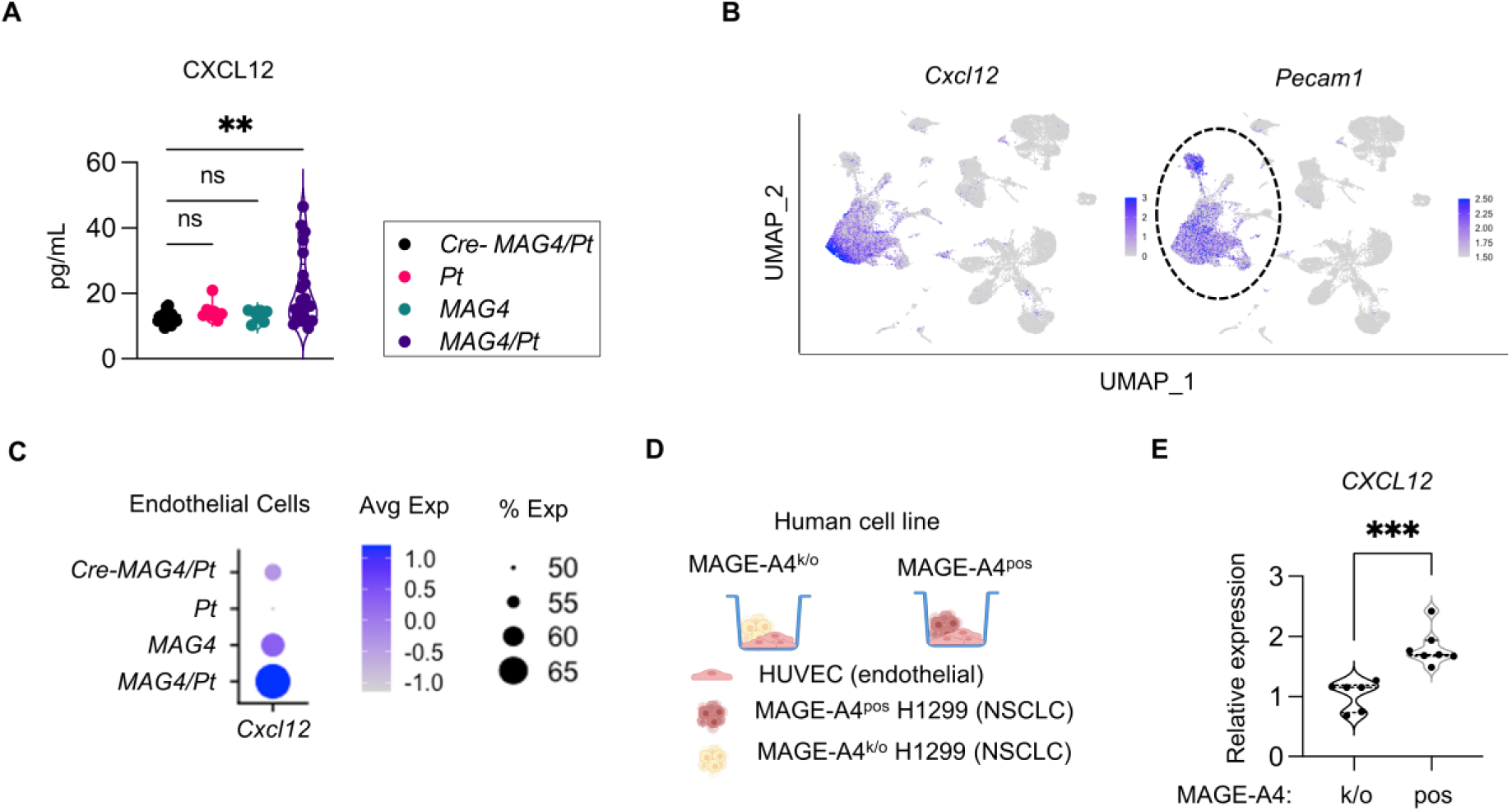
MAGE-A4 induces CXCL12 in endothelial cells to retain plasma cells by CXCL12-CXCR4 axis. **(A)** CXCL12 concentration from whole-lung homogenate from 7-8 months old mice by ELISA. Data from 3 independent experiments combined, n=7-21 per group. (**B**) Featureplot of whole lung shows co-localization of *Cxcl12* with *Pecam1*. (**C**) Dotplot of *Cxcl12* in endothelial (*Pecam1^+^*) cells. (**D**) Schematic of assay design, using Human Umbilical Vein Endothelial Cells (HUVEC) co-cultured with human NSCLC MAGE-A4^k/o^ or MAGE-A4^pos^ H1299 cells for 24hrs. (**E**) *CXCL12* gene expression quantified using qPCR. 2 independent experiments combined, n=3-4 per group. Significance determined by One-way ANOVA with Dunnett’s correction or student t-test. \**P* < 0.05, \*\**P* < 0.01, \*\*\**P* <0.001, ns = not significant.

### MAGE-A4 in human NSCLC shows distinct TIME of increased plasma cells and decreased T cells

To corroborate our findings of the effects of MAGE-A4 on the TIME in human tumors, we used Imaging Mass Cytometry (IMC) with a panel of 38 immune, cancer, and stromal markers (Supplemental Figure 15A, Supplemental Table 2) to assess the immune composition of MAGE-A4^pos^ compared to MAGE-A4^neg^ human NSCLC. We generated a new tissue array using different core samples from 24 of the 29 tumors from Figures 1B and C (Supplemental Table 1). We found that MAGE-A4^pos^ tumors, determined by MAGE-A4 and E-cadherin co-staining (Supplemental Figure 15, B and C), were visibly enriched in CD19^+^ CD38+ CD138^+^ plasma cells (Figure 6, A and B). Quantitatively, CD19^+^ CD138^+^ plasma cells were significantly increased in MAGE-A4^pos^ tumors as total counts and as the relative abundance of infiltrating immune cells (Figure 6, C and D). To assess how MAGE-A4^pos^ tumor sculpts the immediate TIME, we evaluated immune cell density in a 200μm diameter of human NSCLC samples. We found that the plasma cell density in MAGE-A4^pos^ tumor regions was significantly enriched compared with MAGE-A4^neg^ tumors (Figure 6E), suggesting a strong association between MAGE-A4^pos^ tumor and plasma cell localization to tumors. Conversely, the densities of CD8, CD4, and co-localized CD4:CD8 T cells (determined by CD4^+^ CD8^+^ “double-positive” T cells) in MAGE-A4^pos^ tumor were significantly reduced (Figure 6, F-H), demonstrating T cell exclusion and indicating a cold TIME (5, 26). Moreover, density of PD-1^+^ - CD8, -CD4, and -co-localized T cells was decreased in MAGE-A4^pos^ tumor regions (Figure 6, I and J, Supplemental Figure 15D), indicating suppression of T cell activity. Although CD8 T cell counts were unaltered (Supplemental Figure 15, E and G), CD4 T cells and PD-1^+^ CD4 T cells were decreased in MAGE-A4^pos^ NSCLC samples (Supplemental Figure 15, F and H). Similarly, co-localized T cells were decreased, as reflected by the visual absence in UMAP (Figure 6K) and quantitatively as the relative abundance of immune cells (Figure 6L), indicating that interactions between T cells were suppressed in MAGE-A4^pos^ NSCLC (64, 65). Notably, we did not find significantly enriched CD163^+^ CD68^+^ macrophages in MAGE-A4^pos^ tumors (Supplemental Figure 15I). Using immunofluorescent staining, we found that IgA-expressing cells aggregated in regions of MAGE-A4^pos^ tumor in human LUAD and LUSC (Figure 6M, Supplemental Figure 15J). Together, these findings in human MAGE-A4^pos^ NSCLC are consistent with those in *MAG4/Pt* model of murine LUAD and substantiate the role of hMAGE-A4 in promoting the accumulation of IgA plasma cells and fostering a cold TIME in human NSCLC.

**Fig. 6:**
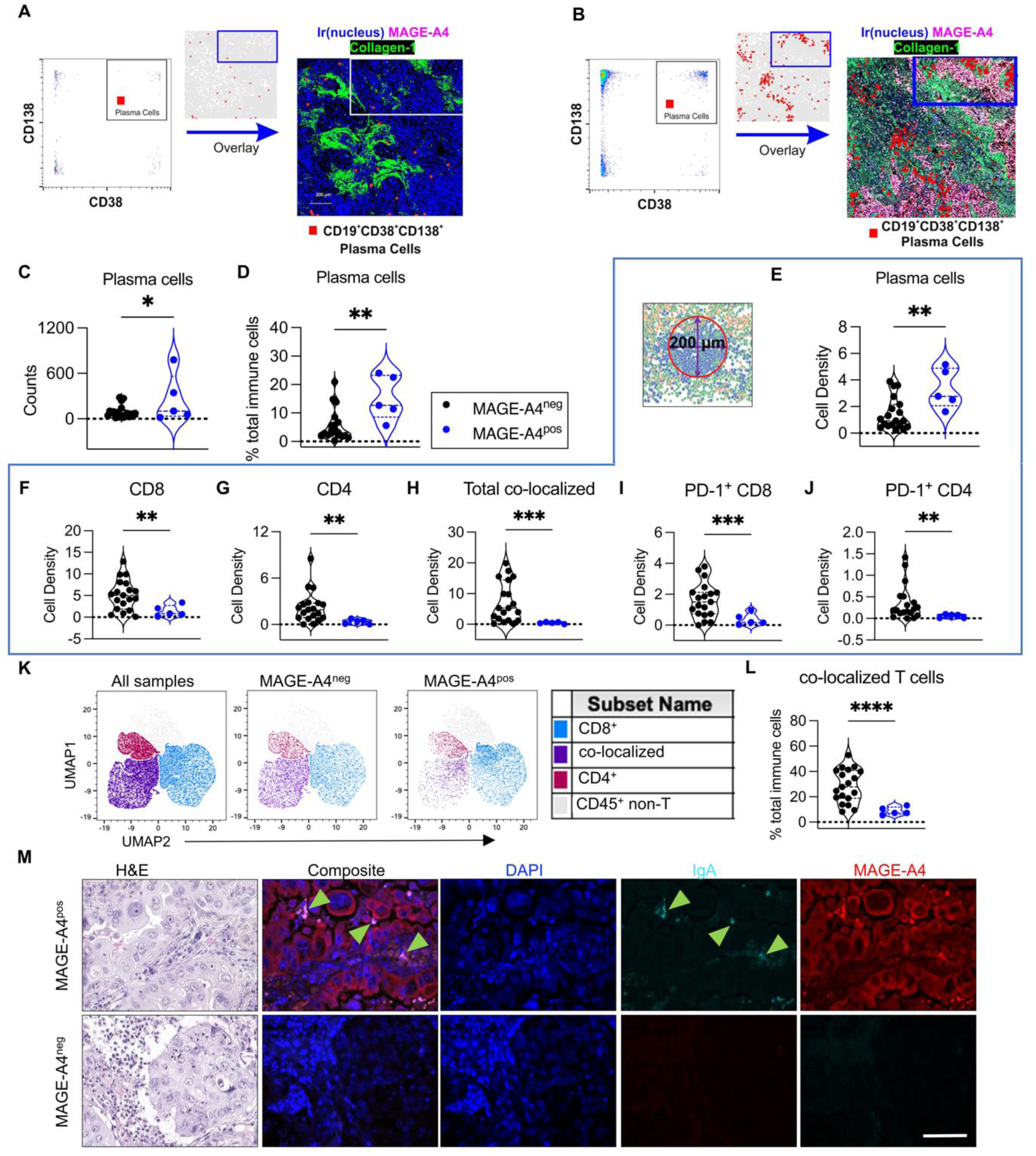
MAGE-A4^pos^ human NSCLC exhibits increased plasma cells and decreased T cells. Imaging Mass Cytometry was utilized to evaluate a microarray of 24 individual NSCLC samples. (**A, B**) Overlay of gated CD19^+^ CD138^+^ CD38^+^ plasma cells onto images showing plasma cells (red) in (**A**) MAGE-A4^neg^ and (**B**) co-localization of MAGE-A4 and E-Cadherin (pink) with plasma cells (red) in MAGE-A4^pos^. (**C**) CD19^+^ CD138^+^ plasma cells quantified as total counts and (**D**) as relative abundance of total immune cells. Total immune cells are the sum of non-B cell CD45^+^, B cells, and plasma cells. (**E-J**) Mean cell density in a 200μm diameter (blue box) around tumor cells (**E**) Cell density calculation schematic (left) and quantification (right) of CD19^+^ CD38^+^ CD138^+^ plasma cells, (**F**) CD8 T cells (**G**) CD4 T cells (**H**) CD8:CD4 co-localized T cells and (**I**) PD-1^+^ CD8 (**J**) PD-1^+^ CD4 T cells. (**K**) UMAP of T cells of all samples and separated by MAGE-A4^neg^ and MAGE-A4^pos^ tumor samples. (**L**) CD8:CD4 co-localized T cells as a relative abundance of immune cells. (**M**) Representative immunofluorescent staining of IgA and MAGE-A4 in lung adenocarcinoma. Green arrows indicate IgA-expressing cells. Scale bar = 75 μm. Significance determined by Welch’s or student t-test. \**P* < 0.05, \*\**P* < 0.01, \*\*\**P* < 0.001, \*\*\*\**P* <0.0001, ns = not significant

### MARPs, but not B220^+^ B cells, drive tumor progression and immune suppression

To determine whether the MARPs prevent or promote tumor growth, we crossed *MAG4/Pt* to *µMt* mice, which lack the IgM heavy chain and fail to develop mature B cells and plasma cells (66), generating B cell and plasma cell deficient *MAG4/Pt* mice (*MAG4/Pt ^µMt^*). We found a significant reduction in tumor burden in 5-month-old *MAG4/Pt ^µMt^* compared to *MAG4/Pt* mouse lung (Figure 7, A and B). Quantification of the tumor showed a ∼60% decrease in tumor burden in *MAG4/Pt ^µMt^* (Figure 7C) and verified a reduction of MARPs (Figure 7D). We next evaluated what immune cells contributed to diminished tumor burden in the absence of B cells and plasma cells in *MAG4/Pt ^µMt^* mice. CD163^+^ TAMs were reduced, CD103^+^ DCs were increased (Figure 7, E and F), and consistently, we found a corresponding increase in Th1 and activated CD25^+^ CD4^+^ T cells (Figure 7, G and H). Notably, PD-1^+^ CD4 and CD8 T cells were enriched in *MAG4/Pt ^µMt^* compared with *MAG4/Pt* mice (Figure 7, I and J). These findings suggest that B cells are critical for tumor development in this model. As the *µMt* mouse has been shown to induce plasmacytoid (p)DCs and high Interferon (IFN)-⍺ in a melanoma model (67), we next assessed B220^+^ CD19^-^ pDCs and IFN-⍺ in *MAG4/Pt ^µMt^* mice. However, we found that neither pDC nor IFN-⍺ were relevant in this model (Supplemental Figure 16A, B). Given that *MAG4/Pt ^µMt^* mice lack both B cells and MARPs (Figure 7D), we next examined the role of B cells in *MAG4/Pt* tumor burden using ⍺-CD20 monoclonal antibody (Supplemental Figure 16C). Compared to isotype control, mice treated with ⍺-CD20 from 2-5 months of age showed significant depletion of B220^+^ B cells but failed to reduce MARPs at 5 months of age (Supplemental Figure 16, D-F). Consistent with the maintenance of MARPs, mice treated with anti-CD20 monoclonal antibodies failed to show any reduction in tumor burden (Supplemental Figure 16, G and H). While an anti-CD20-mediated depletion of B cells reduced the relative abundance of CD163^+^ TAMs, total macrophages remained unchanged (Supplemental Figure 16, I and J), and CD103^+^ DCs did not increase compared to isotype-treated *MAG4/Pt* (Supplemental Figure 16K). These findings suggest that increases in CD163^+^ TAM were driven by B cells, not CD138^+^ plasma cells, while CD103^+^ DCs were inhibited by plasma cells. Nevertheless, the reduction of CD163^+^ TAMs without an increase in CD103^+^ DCs was not sufficient to alter tumor burden, indicating the importance of the antigen-presenting cells in antitumor immunity. Taken together, these data suggest that in this model MARPs, not B220^+^ B cells, mediate tumor progression and the oncogenicity MAGE-A4 in the absence of the tumor suppressor *Pten*.

**Fig. 7:**
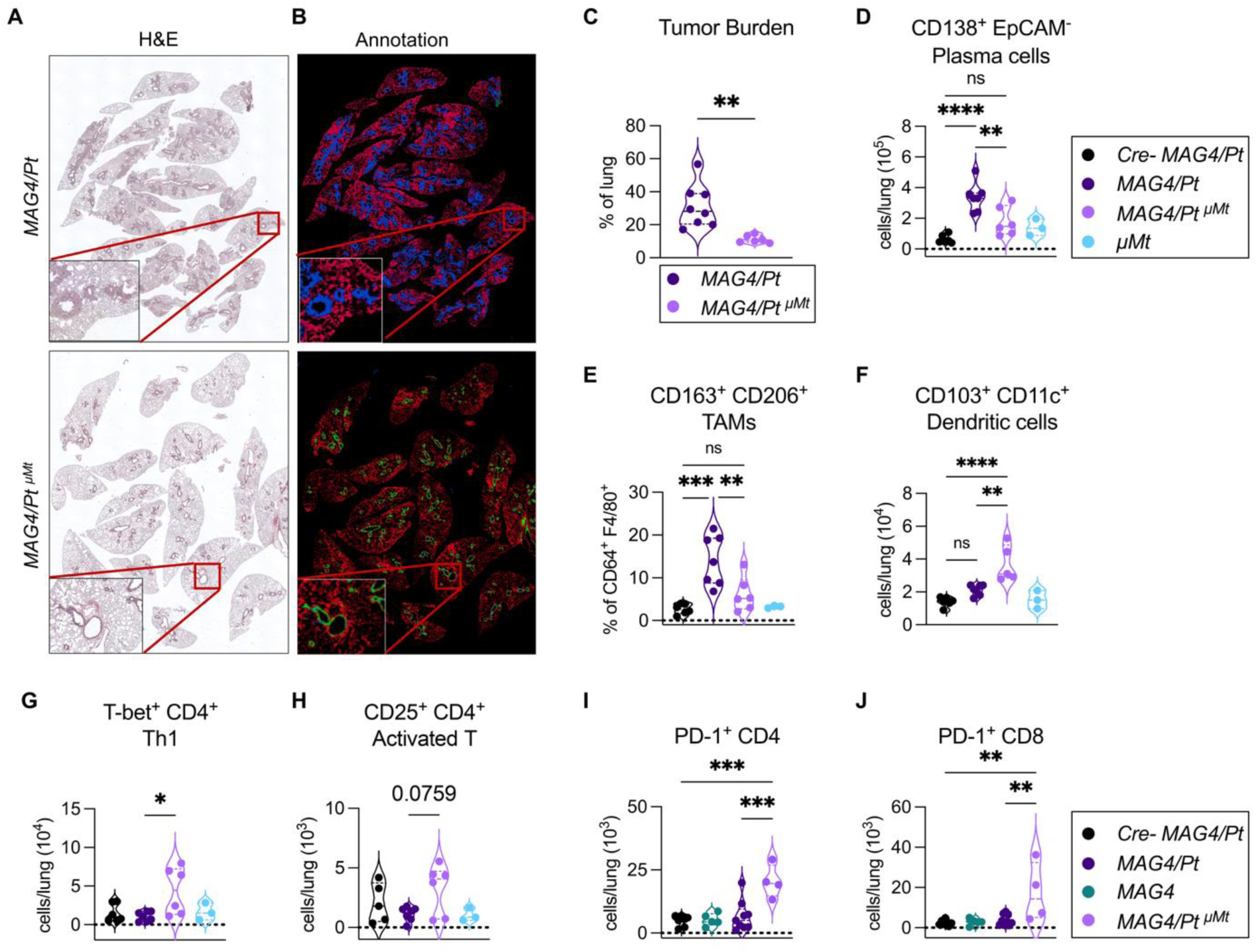
Global B cell ablation reduces *MAG4/Pt* LUAD and restores anti-tumor immunity. (**A**) Representative H&E of *MAG4/Pt* and of *MAG4/Pt μMt* lungs at 5 months of age, 100x. (**B**) Respective annotation from (**A**) of tumor (blue) normal lung parenchyma (red) and normal bronchi (green) using Aivia analysis software. (**C**) Quantification of tumor burden (tumor over total lung area); each data point is the average of 4 measurements per lung, n = 6-7 per group. (**D**-**H**) Flow cytometry of whole lung from 5-month-old *MAG4/Pt μMt* and *μMt* compared with that of historical *Cre-MAG4/Pt* and *MAG4/Pt* mice (**D**) CD138^+^ EpCAM^-^ plasma cells (**E**) CD163^+^ CD206^+^ Tumor-associated macrophages (TAMs) as a percentage of total lung macrophages. (**F**) CD103^+^ dendritic cells (DCs) (**G**) T-bet^+^ CD4^+^ T cells (**H**) Activated CD25^+^ CD4^+^ T cells. (**I**) Total lung PD-1^+^ CD4 and (**J**) PD-1^+^ CD8 T cells from 5-month-old mice; representative of 2 independent experiments, n= 4-9 per group. Significance determined by One-way ANOVA with Tukey’s correction or student t-test. \**P* < 0.05, \*\**P* < 0.01, \*\*\**P* <0.001, \*\*\*\**P* <0.0001, ns = not significant.

## Discussion

Human lung cancer is characterized by an altered TIME and genetic aberrations, including mutations and expression of CTAs (1, 68–70). We show that while expression of hMAGE-A4 in the airway epithelia alone is not sufficient for cancer development, its expression synergizes with airway-specific *Pten* loss to induce lung adenocarcinoma. Tumors expressing hMAGE-A4 are invasive, develop as early as 3 months of age, and metastasize rapidly, demonstrating that this CTA accelerates the development and progression of cancer in susceptible hosts. The expression of MAGE-A4 has been shown to inhibit apoptosis (71) and prevent accurate DNA repair mechanisms (72), both of which could impact additional cell damage resulting from exposure to carcinogens such as those in cigarette smoke (73). Our findings here support the role of hMAGE-A4 in the inhibition of antitumor immunity and accumulation of immunosuppressive MARPs (Graphical Abstract).

Human clinical studies suggest a strong association with MAGE-A4 expression in LUSC, however, the expression of this CTA in the lungs in the absence of *Pten* resulted in adenocarcinoma in mice. There is some precedent for genetic markers in human tumors that do not closely follow the expected histopathology in mice, suggesting that animal models serve as an approximation of human diseases (74, 75). Specifically, signaling pathways are known to differ between mice and humans in cis-regulatory elements (e.g., P53) that target transcription factors binding large areas of DNA, suggesting that multiple species-specific factors play a role in malignant transformation and histological phenotypes (74).

Consistent with a cold tumor phenotype observed in human NSCLC subtypes (76–78), we found reduced T cell infiltration in *MAG4/Pt* lung. Indeed, neither pro- nor antitumor T cells were induced, as evidenced by the lack of activation or inhibitory markers such as PD-1 and the lack of Tregs. The paucity of T cells and the absence of activation, inhibitory, or exhaustion markers indicates a deficiency of response and recognition of tumors by T cells. This observation has several implications with clear relevance in clinical applications. For example, the potential use of affinity-optimized T cells targeting MAGE-A4 antigen to NSCLC (13) should be assessed against the profound immunosuppressive effects of MAGE-A4 that inhibit T cell trafficking to the tumor, possibly mitigating antitumor effects of therapeutic T cells. Further, ablation of CD138^+^ plasma cells resulted in increased PD-1^+^ CD8 and PD-1^+^ CD4 T cells in the lungs. These findings reveal that targeting MARPs may be an avenue to polarize MAGE-A4^pos^ NSCLC to increase T cell infiltration and theoretically sensitize NSCLC to immune checkpoint blockade. Together, these findings unveil a potential mechanism to polarize human MAGE-A4^pos^ NSCLC from a cold tumor microenvironment to one that enables T cell infiltration and optimizes antitumor immunity. Whether plasma cell depletion improves response to immune checkpoint blockade in MAGE-A4^pos^ NSCLC warrants evaluation in future studies.

Macrophages have been associated with anti- or pro-tumor effector functions, the latter of which reside in cold tumor microenvironments (79). While we did not observe a significant increase in CD163^+^ TAMs in our human IMC, larger dataset analyses have revealed that CD163^+^ TAMs are increased in MAGE-A4^pos^ human NSCLC (80), validating the role of MAGE-A4 in driving this subset and promoting a cold TIME. Furthermore, CD163^+^ TAMs were expanded in *MAG4/Pt* lung and were reduced after anti-CD20-mediated B cell depletion, which was not sufficient to alter tumor burden, indicating that CD163^+^ TAMs were not a main immune contributor to tumor progression in MAGE-A4-driven NSCLC. However, total macrophages and antitumor CD103^+^ DCs were increased in *MAG4/Pt ^µMt^* mice in addition to reduced TAMs. Increased DCs likely contribute to the increase in T cell infiltration and activation, as well as to the reduction in tumor burden seen in *MAG4/Pt ^µMt^* compared with *MAG4/Pt*.

In human non-EGFR LUAD, expanded plasma cells correlate with decreased memory B cells (81). Accordingly, we showed that memory B cells were not enriched, while plasma cells were increased in *MAG4/Pt* lungs, indicating the alignment of this model with the immunological characteristics of human NSCLC. A retrospective study of human LUAD showed that while the presence of CD20^+^ B cells is beneficial, a higher ratio of plasma cells to total infiltrating lymphocytes was associated with reduced survival (40). Similarly, in *MAG4/Pt* lung, an increase in MARPs without a concomitant increase in B or T cells in the lungs promoted tumor progression. Furthermore, we show that plasma cells, not B cells, promote tumor development, as depletion of B cells by anti-CD20 did not affect tumor burden, but global B- and plasma cell ablation (*MAG4/Pt ^μMt^*) decreased total tumor in the lungs.

Plasma cells enriched in human LUAD predominantly express IgA (81). A high IgA:IgG ratio has also been associated with lower survival in patients with NSCLC (81, 82), and IgA in other tumors has been implicated in an immunosuppressive and pro-tumor function (31). Similarly, the pro-tumorigenic plasma cells observed in *MAG4/Pt* tumors produce IgA and the cytokines IL-10 and TGF-β, both of which have potent immunosuppressive capacities (32, 83, 84). While it has been similarly observed that IgA-producing plasma cells also express IL-10 (48), here we used IgA expression as a marker of a specifically polarized plasma cell; future studies are warranted to assess the specific function of IgA in NSCLC progression and immunosuppression.

Tumors with infiltrating B cells have increased CXCL13 (60, 81, 85). We did not observe any increase in CXCL13 protein in lung homogenate supernatant, consistent with the lack of B220^+^ B cell infiltration. However, the expression of CXCL12, a chemotactic factor for plasma cells, (59) was increased in *MAG4/Pt* lung. The CXCL12-CXCR4 axis is a well-established mechanism of plasma cell retention in the bone marrow (59). We show that *MAG4* and *MAG4/Pt* endothelial cells have upregulated *Cxcl12* and that MAGE-A4^pos^ cancer cells induce human endothelial cells to upregulate the expression of *CXCL12*. Thus, CXCL12 likely contributes to the retention of MARPs in *MAG4/Pt* lung, a finding that has significant clinical implications in the treatment of patients with MAGE-A4^pos^ NSCLC. Further, because MAGE-A4 is expressed in many solid tumors, whether our findings extend beyond NSCLC should be evaluated in future studies.

Our findings that hMAGE-A4 drives tumorigenesis and substantially impacts the tumor immune microenvironment are of clinical importance considering the effort that has been invested in targeting MAGE-A4. Consistently, targeting MAGE-A4 in various cancers using immune-mediated methods (86–88) and autologous T cells against MAGE-A4 in Phase I Clinical trials has shown some promise (13). However, the findings here could impact tailoring new treatments in MAGE-A4^pos^ solid tumors. Given that our work has revealed that MAGE-A4 suppresses T cell responses, we anticipate that these T cell therapies may be inhibited by the immune microenvironment created by MAGE-A4-driven tumors and warrants further investigation in clinical studies. Depleting plasma cells could potentially be used in combination with MAGE-A4 targeted therapies to increase infused T cell infiltration and activity. Checkpoint blockade could also be combined with plasma cell depletion methods, as we have shown that in *MAG4/Pt ^µMt^* lung, the immune microenvironment is remodeled and PD-1^+^ T cells emerge. Our work reveals not only the role of MAGE-A4 in driving tumor progression but also a targetable, immune-mediated mechanism.

Collectively, these findings reveal additional questions and work needed to further understand the immunological and cancer intrinsic effects of MAGE-A4. Our studies used clinical observations (e.g., the association between hMAGE-A4 expression and loss of *Pten*) to develop a relevant model of NSCLC. However, other genetic susceptibilities associated with MAGE-A4 may drive tumor progression through immunosuppressive plasma cells. MAGE-A4 is expressed in many other types of cancers, including melanoma, head, and neck, ovarian, gastric, and others (89, 90), thus its effect may not be restricted to the lung tissue. Genetic background or organ-specific context may alter the immune response or contribution of MAGE-A4. Regardless, our findings in human MAGE-A4^pos^ NSCLC corroborate its effects in mice and humans. Therapeutically, our study is limited to the evaluation of plasma cells by global knockout, rather than by specific depletion. Various plasma cell depletion modalities could be further investigated (91, 92) in MAGE-A4^pos^ NSCLC.

In summary, our findings demonstrate for the first time that MAGE-A4 has tumorigenic potential and significantly alters the tumor immune microenvironment. MAGE-A4 elicits tumor progression by promoting the accumulation of MARPs which suppress macrophage, dendritic cell, and T cell infiltration and activation, while promoting a cold immune microenvironment which results in tumor progression and metastasis. This study further highlights a potential therapeutic vulnerability and new possibility in restoring antitumor myeloid and T cell immunity.

## Methods

### Mice

To account for sex as a biological variable, both male and female mice were used in all experiments in approximately equal proportion, and no sex-biased outcomes were observed. Thus, all the data is combined. The exception is scRNA-seq experiments, where only female mice were used, as only one sample per genotype was run. However, even in these experiments, the questions we addressed and the results obtained were relevant to both sexes, specifically changes of the tumor microenvironment, which we validated in further experiments.

The Shuttle Vector (RfNLIII), base vector (pCAGGS-LSL-luciferase), SW102 cells for homologous recombination by electroporation, flp recombinase in the 294-FLP bacterial strain were generously provided by Dr. Ming-Jer Tsai (formerly at Baylor College of Medicine). To generate an embryonic stem cell targeting construct, the human MAGE-A4 cDNA (Origene, RC223991) was inserted into the targeting construct downstream of a Lox-Stop-Lox (LSL) cassette (Supplemental Figure 2A). Targeted ES clones were identified by Southern analysis of their EcoRV digested genomic DNA. Positive ES clones, which were heterozygous for the targeted event, were scored by the presence of both 11.5 and 3.8 kb hybridization bands corresponding to the wild-type and targeted Rosa26 alleles respectively (Supplemental Figure 2B). The positive embryonic stem cells were injected into albino C57BL/6N strain blastocysts and transferred them to female recipients to generate chimeras. Standard procedures were used to generate chimeric founder (F0) mice from targeted ES clones (93, 94). A minimum of three individual targeted ES clones were used to generate separate monogenic *MAGE-A4^LSL^* mouse lines, each of which harbored a single copy of the minigene. Chimeras were bred with C57BL/6 mice. For expression of human MAGE-A4, *MAGE-A4^LSL^* mice were crossed with the club cell specific *CCSP^iCre^* transgenic mouse (42, 95). Resultant *CCSP^iCre^: MAGE-A4^LSL^* bigenic mice are referred to as MAG4 mice. *MAGE-A4^LSL^: Pten^f/f^* was crossed with *CCSP^iCre^:MAGE-A4^LSL^* to generate the trigenic *CCSP^iCre^: MAGE-A4^LSL^: Pten^f/f^* (*MAG4/Pt*) mouse. Mice were maintained on a C57/B6 background. For metastasis screening studies, *MAG4/Pt* mice were further crossed to Luciferase-expressing mice to result in MAGE-A4-heterozygous and Luciferase-heterozygous mice. ROSA26-pCAGGs-LSL-Luciferase mice were a gift from Dr. Ming-Jer Tsai, originally obtained from NCI. Wild-type, *μMt*, and *Pten^f/f^* mice (all C57BL/6 background) were originally purchased from the Jackson Laboratory. *CCSP^iCre^* (C57BL/6) mice were acquired from Dr. Franco DeMayo’s laboratory at Baylor College of Medicine.

### In silico data

For evaluation of MAGE-A4 expression delineated by tumor stage, a subset of The Cancer Genome Atlas (TCGA) data was used as curated by the Human Protein Atlas (accessed Dec. 2022); a total of 989 NSCLC samples were assessed. Fragments per kilobase million (FPKM) of greater than or equal to 1.0 was considered positive. For correlation analysis of MAGE-A4 expression with loss of tumor suppressors, a total of 1023 NSCLC cases from TCGA and 337 genomics cases from CPTAC datasets were analyzed as previously described (9). Briefly, RNA-seq data were obtained from the Broad Institute Firehose pipeline (http://gdac.broadinstitute.org/).

Copy-number alteration data and somatic mutation data were obtained (41). For TCGA copy number alteration data, we obtained SNP array-based CNA “threshold” values (−2, −1, 0, 1, 2) from the Broad Institute’s Firehose data portal [https://gdac.broadinstitute.org]. For CPTAC data, copy number data were first obtained from the Genome Data Commons [https://gdc.cancer.gov/]. We first normalized the absolute copy data according to ploidy (dividing gene copy value by average copy value for all genes), then threshold to values approximating homozygous deletion (−2), heterozygous deletion (−1), wild-type (0), gain of 1–2 copies (+1), and amplification with at least 5 copies (2). It has been previously shown that even single copy losses of PTEN have a notable impact on PTEN expression and downstream pathway signaling (96, 97), so single copy losses were included in loss of PTEN. For heat map generation and for gene signature scoring, gene expression values were log-transformed and then z-normalized to standard deviations from the median across all cancer samples. JavaTreeview (version 1.1.6r4) was used for heatmap visualization (98).

### Flow cytometry

Lungs were perfused, chopped into ∼1mm pieces, then digested using 0.15mg/mL of Liberase TL (Sigma, 5401020001) and 0.04mg/mL of DNAse (Sigma, 10104159001), in RPMI (Gibco) for 25 min at 37 °C. Tissue was then passed through a 70μm filter and erythrocyte lysis was performed using in-house Ammonium-Chloride-Potassium (ACK) lysing buffer. Cells were blocked for Fc receptors using Truestain FcX plus (Biolegend, 156604,S17011E) then stained for specified surface-markers. Flow cytometry antibodies used are in Supplemental Table 2. Cell viability was evaluated using LIVE/DEAD fixable blue dead cell stain kit (Thermo Fisher, L23105). For intracellular cytokines, single-cell suspensions were stimulated with phorbol 12-myristate 13-acetate (PMA) (10 ng/ml) (Sigma) and ionomycin (250 ng/ml) (Sigma) supplemented with brefeldin A (10 µg/ml) (Sigma) for 4 hours. After surface staining, cells were fixed using 2% PFA, or fixed and permeabilized using the Cytofix/Cytoperm Fixation/Permeabilization Solution Kit for intracellular markers (BD Biosciences) or the Foxp3/Transcription Factor Staining Buffer Set (Thermo Fisher) for transcription factors, each according to manufacturer protocols. Cells were analyzed on an LSRII or sorted on FACSAria Fusion with FACSDiva software (BD Biosciences) and analyzed with FlowJo software (TreeStar).

### Lung Histology, Immunohistochemistry (IHC), Immunofluorescence (IF), and Imaging Mass Cytometry (IMC)

Twenty-nine paraffin-embedded de-identified NSCLC cases were obtained from the Michael E. DeBakey Veteran’s Affairs Medical Center in Houston, TX and placed to create a tumor array. 5 μm sections were deparaffinized and rehydrated for IHC or IMC staining. For IMC staining, the protocol provided by Standard Biotools was used and antibodies used are found in Supplemental Table 3. One-mm^2^ region of interest (ROI) was acquired per sample in the Hyperion Imaging System (Fluidigm). For mice, freshly harvested tissues were fixed in 4% paraformaldehyde (PFA) overnight before being processed and embedded in paraffin according to standard protocols. Hematoxylin and eosin (H&E), IHC, and IF staining for either human arrays or mouse tissue were performed as previously described (42) using Diva Decloaker (BioCare Medical) for heat-induced antigen retrieval and antibodies against targets listed in Supplemental Table 4. Images of immunostained tissue sections were captured using Evos Microscope (ThermoFisher). Post image processing was performed using Adobe Photoshop^®^ software programs (Adobe Systems Inc.) and ImageJ (NIH). For IMC analysis, the quality of the samples was evaluated by inspecting marker-staining patterns within the HistoCAT2 Viewer (https://github.com/BodenmillerGroup/histoCAT3D) (99). Raw data, stored as .mcd files, were transformed into TIFF format using a specialized Python package adapted from the ElementoLab IMC Python Package (https://github.com/ElementoLab/imc). These TIFF files were further analyzed with the Ilastik tool to predict cellular locations and boundaries via pixel and object classification (100). Single-cell segmentation for each image was achieved using the DeepCell package (101, 102). Nuclear analysis was integrated with cell boundary data for cell segmentation. The mean expressions of panels from segmented single cells were retrieved using the IMC package by overlaying segmentation masks onto the corresponding TIFF images. Resulting segmented images and data files were catalogued into CSV and AnnData formats. For enhanced precision in cell protein expression metrics, we employed the PowerTransform (Yeo-Johnson) normalization tools designed for skewed and bimodal condition interpretations. This normalization was executed for each region of interest (ROI; each ROI was an individual sample). For cell-type identification, a cell-type tree diagram was gated with the FlowJo™ v10 Software (BD Life Sciences). To evaluate cell-cell interactions, we employed a permutation test approach available in the SCANPY and SCIMAP packages. This method determines if interactions or avoidances within or between cell types surpass random occurrence frequencies (103). The IMC images were transposed into topological neighborhood graphs, wherein cells were depicted as nodes. Direct cell-cell neighboring pairs, within a 200µm range between centroids, were depicted as edges. The quantification of neighborhoods is predicated upon the cellular density surrounding the cancer, facilitating comparisons between grouped data sets. Confirmatory visualizations underpinned the results, overlaying original images with specific cellular compositions.

### RNA extraction and real-time quantitative PCR

Cells were isolated using the RNeasy^®^ Plus Mini Kit (Qiagen Inc.) according to manufacturer’s protocol. Splenic tissue was treated with TRIzol (Invitrogen), and mRNA was extracted by phenol/chloroform (Sigma-Aldrich). cDNA was made with the High-Capacity cDNA Reverse Transcription Kit (Applied Biosystems) and RT-qPCR was performed using the ABI PerkinElmer Prism 7500 Sequence Detection System (Applied Biosystems) using Taqman probes *Scgb1a1* Mm00442046_m1, *CXCL12* Hs00171022 (ThermoFisher), *18S* ribosomal RNA (Hs99999901_s1) and Taqman Universal PCR Master Mix (Applied Biosystems).

### Immunoblotting

Western immunoblotting conditions have been reported previously (104). Primary antibodies are provided in Supplemental Table 4.

### scRNA-seq

Whole lung from all four genotypes were used for single-cell sequencing; 1-3 lungs per sample were used and pooled into one sample per genotype for 10X sequencing. Single cells were obtained from lung by perfusing the left ventricle with ice-cold PBS then placing in digestion media prepared from the Lung Dissociation Kit (Miltenyi) and digested on a gentleMACS Octo Dissociator, using pre-set program 37_m_LDK_1 (Miltenyi). Cells were passed through 70 μm filter and treated with ACK for red blood cell lysis for 2 min and stained with DAPI to sort live cells on a Sony SH800 sorter using a 70 μm sorting chip. Cells were processed, barcoded, and analyzed on 10X genomics platform per the 10X genomics protocol. Raw sequencing files were imported into the 10x Genomics Cell Ranger toolkit (v3.1.0) for alignment, filtering, barcode counting, and UMI counting with default parameters. The mm10 (v3.0.0) genome was used for reads mapping. The Seurat (v3.2.3) package (105, 106) on R (v4.0.2) was used for downstream analysis. For quality control, we kept cells with fewer than 20,000 read counts and less than 10% mitochondria genes to filter doublets and dead cells. We removed mitochondrial genes (genes start with mt-), ribosomal protein genes (genes start with Rpl and Rps), and unspecified genes (genes start with Gm) from analysis. The *SCTransform* function (107) in the Seurat package was used to implement regularized negative binomial regression to normalize raw UMI counts. Variable genes identified by this method were used as inputs for principal component analysis (PCA). We applied Uniform Manifold Approximation and Projection (UMAP) analysis for visualization using the first 50 PCs identified by PCA. Cells were clustered by the shared nearest neighbor (SNN) with the *FindNeighbors* and *FindClusters* functions in the *Seurat* package. SingleR (52) was used to identify cell cluster identities, which were validated by common cell identity markers.

### Tumor burden studies and analysis

To evaluate tumor burden, lungs were sliced into 1-2 mm portions along the transverse plane and embedded in paraffin to maximize the area of the lung represented in any section. For ⍺-CD20 depletion experiment, only the left lobe was evaluated; otherwise, the entire lung was used. Blocks were then sectioned through the entire lung, taking 5 μm sections every 100-150 μm. Of these, 4 sections representing similar depths of the lung across samples (approximately every 400-600 μm) were H&E stained and evaluated for tumor burden. Sections were imaged with Leica microscope at 100x, with multiple images acquired using MicroManager from ImageJ (NIH), processed to remove background in FIJI, and stitched in Aivia software (Leica) for full representation of the entire tissue block. Aivia was used to annotate and train the classifier to recognize the categories: background, tumor, normal lung parenchyma, or normal bronchi. The trained classifier was applied to all the full, stitched images. Tumor burden was calculated as area of tumor divided by area of total lung (tumor burden = tumor / (tumor + lung parenchyma + normal bronchi)); for each individual lung, tumor burden was reported as an average of the 4 analyzed sections to result in one data point per mouse.

⍺-CD20 antibody (5D2) or isotype control (GP120:9674) (Genentech) were given at 200μg once per week for 3 weeks beginning at 2 months of age, then every other week thereafter until 5 months of age, at which time the lungs were perfused and harvested: the left lobe was taken for histology and the right lobes were processed for flow cytometric analyses. Metastatic potential was assessed in *MAG4/Pt Luc* mice by bioluminescence imaging using the In-Vivo MS FX Pro system (Bruker Corporation). 2-3 minutes after 150mg/kg D-luciferin (ProMega) intraperitoneal administration to allow for circulation, mice were euthanized, organs removed, and imaged within 10 min of D-luciferin administration.

### ELISAs and LEGENDPlex

Lung homogenate supernatant, bronchoalveolar lavage (BAL) fluid, or serum was used for sample as indicated. Lung homogenate supernatant was collected from whole lung digested as described above for flow cytometric analyses. Cells were cultured in complete medium for 16-18 hours, then the supernatant was collected. BAL was collected by intratracheal perfusion of 800 μL of ice-cold PBS twice for a total of 1600 μL, then collecting the cell-free supernatant. ELISAs were used per kit directions CXCL12 and CXCL13 (DuoSet ELISA kit, R&D Systems), and IL-10 (BD Biosciences). Immunoglobulin Isotyping Mouse Uncoated ELISA kit (ThermoFisher, 88-50630-86) and isotype control antibodies were used as standards (Supplemental Table 4). LEGENDPlex 13-plex mouse-antivirus response panel (Biolegend, 740621) and provided Qognit software was used to analyze IFN-⍺.

### In vitro cell experiments

H1299 cells were a gift from Dr. Edwin Ostrin (MDACC) and verified by short tandem repeat (STR) profiling at the Cytogenetics and Cell Authentication Core at MDACC. Cell cultures were not verified to be mycoplasma free. MAGE-A4 knock-out clones were generated from H1299 cells using CRISPR plasmid and control scramble plasmid (SantaCruz, sc-403486 and sc-418922) and Lipofectamine 3000 (ThermoFisher), following manufacturer protocol specific for H1299 cells. After transfection, single cells expressing GFP were isolated by fluorescence using an Aria cell sorter into a 96-well plate and subsequently expanded. Clones were verified for MAGE-A4 knockout by western blot (sc-294929, Santa Cruz Biotechnology). Pooled HUVECs (ATCC) were expanded in Vascular Cell Basal Media per manufacturer protocol and seeded at a density of 2x10^5^ cells; H1299 cells seeded at 1x10^5^ cells, total volume of 800μL in a 12-well plate and cultured for 24 hr in F12/DMEM media supplemented with 10% FBS, 5% L-glutamine, and 5% Pen/Strep (Gibco). HUVECs alone were seeded on plates coated with 0.01 mg/mL fibronectin, 0.03 mg/mL bovine collagen type I and 0.01 mg/mL bovine serum albumin. Cells were treated with recombinant MAGE-A4 (Abcam, ab99138), vehicle (dimethyl sulfoxide, DMSO), or PX-478 (MedChemExpress, HY-10231) as indicated in figure legends. Cells were lysed and RNA extracted using RNeasy Mini Kit (Qiagen) according to manufacturer protocol.

### Statistics

Statistical analysis and graphs were generated using GraphPad Prism 10 software (GraphPad). Data are presented as individual data points and visualized in violin plots. Statistical comparison between groups was performed using the unpaired Student’s t-test or One-way ANOVA with Tukey’s or Dunnett’s correction as appropriate. Occasionally, outlier test was performed in flow cytometry results using ROUT’s test and resulting outliers removed from final analysis. For IMC data analysis, Levene’s test was applied; where F≥4, Welch’s t-test was used; otherwise, student t-test was applied. A *P*-value less than 0.05 was considered statistically significant, ns indicates not significant. *P* value and *N* are noted in figure legends.

### Study approval

All mice were bred, housed, and/or treated in the transgenic animal facility in accordance with the guidelines of the Institutional Animal Care and Use Committee at Baylor College of Medicine. Human lung samples were used by consent under approval from the Institution’s Review Board protocol H-10797 at the Michael E. DeBakey Veteran’s Affairs Medical Center.

### Data availability

scRNA-seq data will be deposited to the Gene Expression Omnibus (GEO) database upon acceptance of manuscript. All data is included in manuscript and values for all data points in graphs are reported in the Supporting Data Values file. Analytic code is available as supplemental file.

### Author contributions

DA, CYC, FK contributed to research design, conceptualization, methodology, data interpretation, and visualization. DA, CYC, MJH, SJ, BC, AC, and LZS performed experiments. DA, CYC, AC, LG, YS, CJC, AC, NJK, SWK, WH, and HSL analyzed and interpreted data. DA wrote the original draft; all authors contributed to the review and editing of the manuscript.

## Acknowledgements

This project was supported by the Cytometry and Cell Sorting Core at Baylor College of Medicine (BCM) with funding from the Cancer Prevention & Research Institute of Texas (CPRIT) Core Facility Support Award (CPRIT-RP180672), the National Institutes of Health (NIH) (CA125123 and RR024574) and the expert assistance of Joel M. Sederstrom and by the Mouse Metabolism and Phenotyping Core at Baylor College of Medicine, subsidized through BCM’s Advanced Technology Cores and NIH funding (UM1HG006348, R01DK114356). We also thank Sang Jun Han, Sung-Nam Cho, and Mark S. Kim for their technical support and Texas Children’s Hospital William T. Shearer Center for Human Immunobiology for their generous support of this research. Figures 5D and S17 were created in BioRender.com. This work was supported by NIH grants: R01 ES029442-01, R01 AI135803-01, and the US Veterans Affairs (VA) Merit grant CX000104, VA I50CU000161; DOD W81XWH-20-1-0607 to F.K.; DLDCCC pilot award to FK and WH; NIH grants HL117181, HL140398, R01 AI135803, and R41 AI124997 and VA Office of Research and Development grant I01BX004828 to D.C., DOD W81XWH-22-1-0657 and CPRIT RP200443 to HSL, and NIH T32GM088129 and Baylor Research Advocates for Student Scientists (BRASS) scholarship to D.A. The content is solely the responsibility of the authors and does not necessarily represent the official views of the NIH.

