## Supplemental for "MAGE-A4-Responsive Plasma Cells Promote Non-Small Cell Lung Cancer"

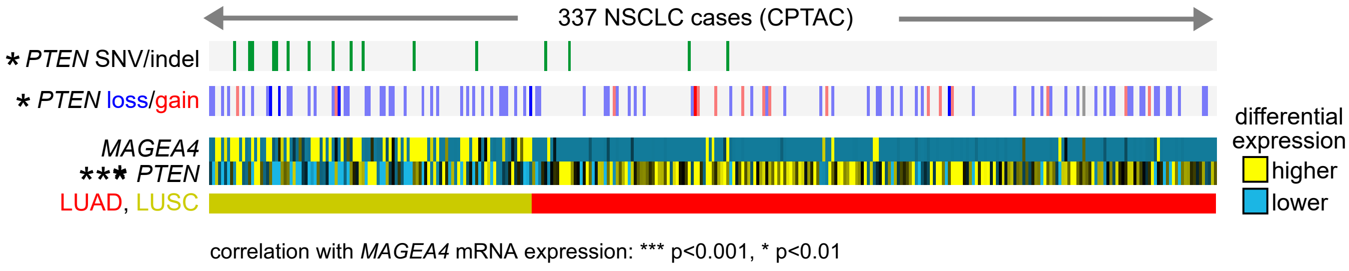

**Fig. S1: MAGE-A4 expressed in human NSCLC is associated with loss of *PTEN* in an independent cohort**. Analysis of MAGE-A4 mRNA expression in human NSCLC correlated with decreased *PTEN* copy and mRNA levels using data from the Clinical Proteomic Tumor Analysis Consortium (CTPAC) (N= 337). Association of MAGE-A4 with *PTEN* mRNA and copy number levels by Pearson's correlation.

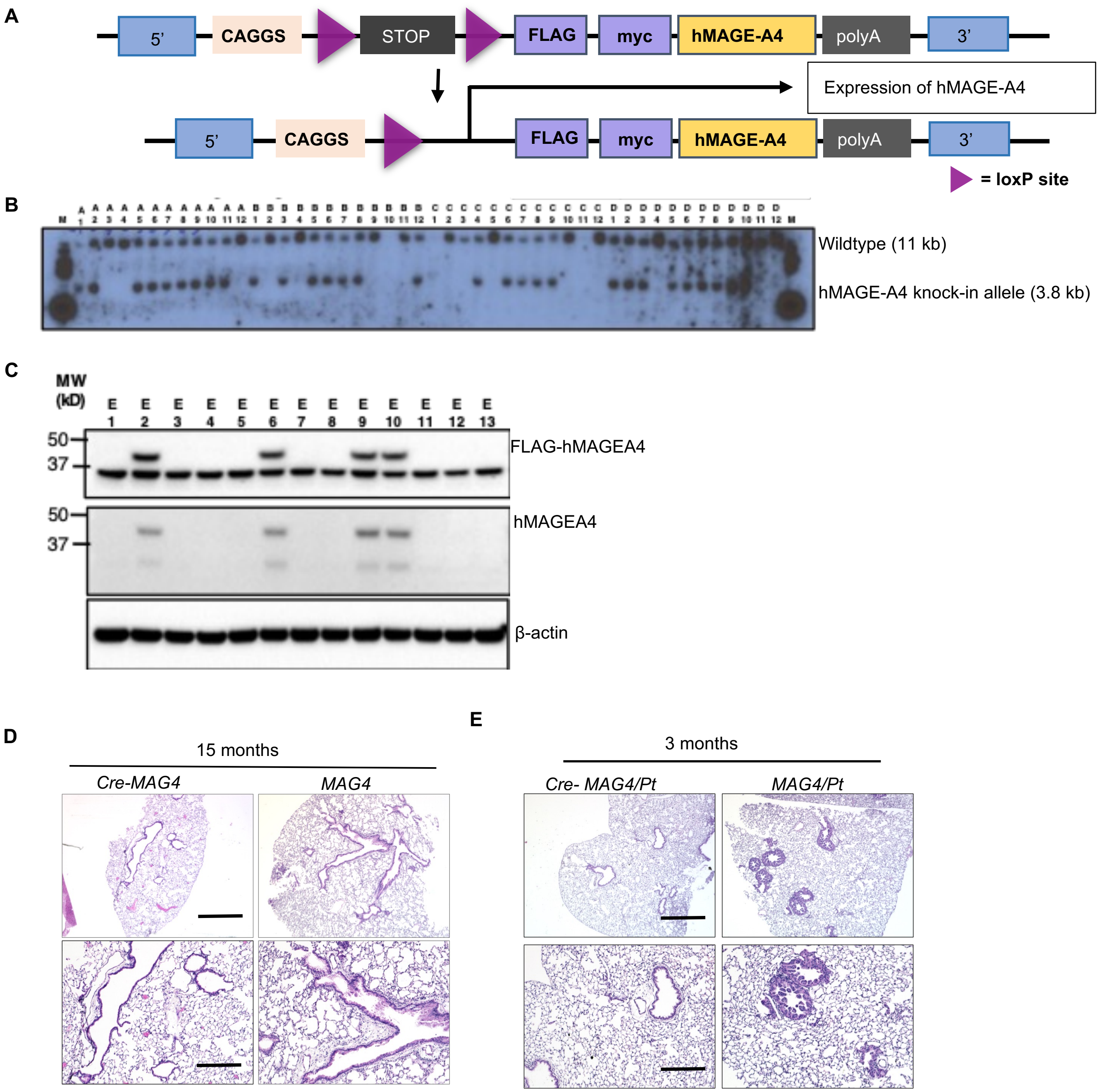

**Fig. S2: Generation of expression of MAGE-A4 in mouse model and resulting phenotype**. (**A**) Diagram of expression minigene and method of MAGE-A4 expression. The targeting vector contains 5' and 3' arms of Rosa26 genomic sequences, flank a minigene, which consists of (in the 5' to 3' direction): the CAGGS promoter; a LoxP-STOP-LoxP (LSL) cassette; a cDNA encoding the human Melanoma-associated antigen A4 (MAGE-A4); and a polyadenylation signal. (**B**) Southern blot to verify embryo h*MAGE-A4* knock-in. (**C**) Western-blot verification of MAGE-A4 expression and FLAG-tag expression in whole lung protein isolate. (**D**) *MAG4* representative lung histology (H&E) at 15 months of age. Scale bars 75 µm (top) and 300 µm (bottom). (**E**) Representative image of tumor in *MAG4/Pt* at 3 months of age (H&E) Scale bars 75 µm (top) and 300 µm (bottom).

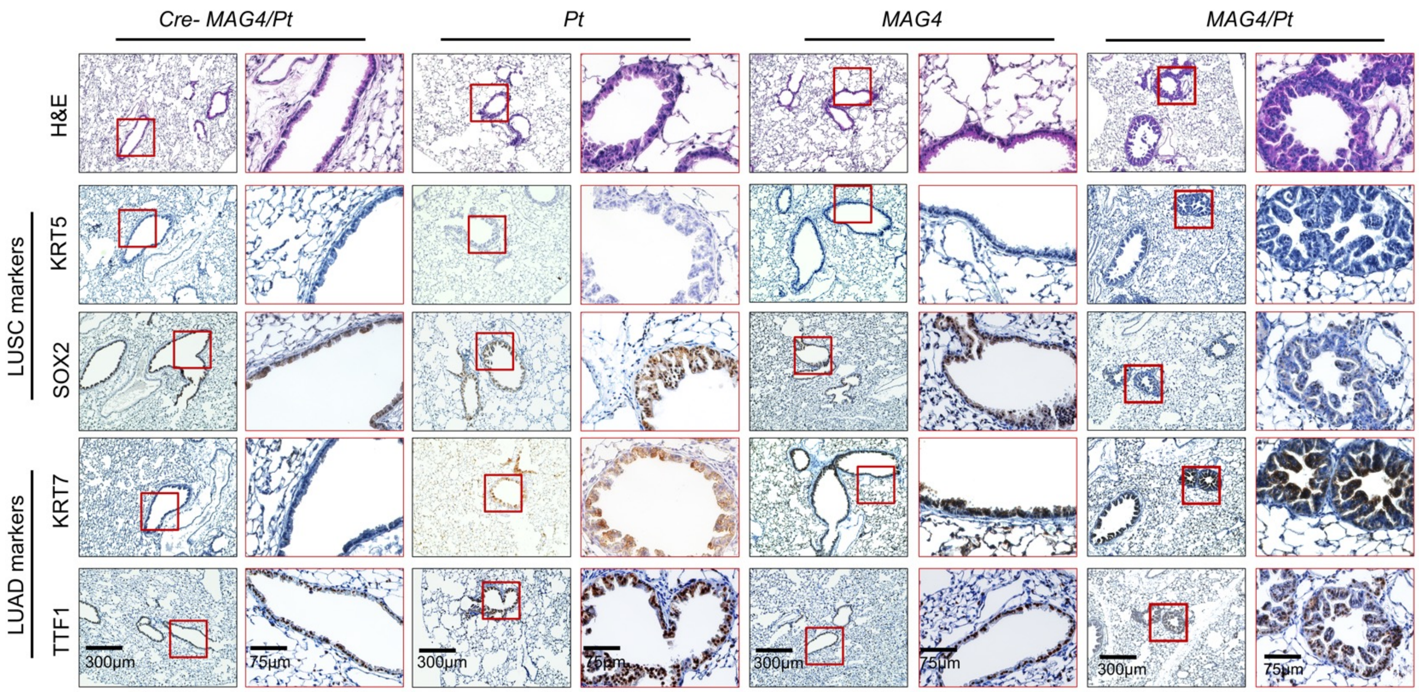

**Fig. S3: *MAG4/Pt* tumors are invasive and are negative for squamous cell but positive for adenocarcinoma**. Immunohistochemical staining for markers of squamous cell (LUSC) (i.e. KRT5 and SOX2) and adenocarcinoma (LUAD) (i.e. KRT7, TTF1) in *Cre-MAG4/Pt*, *Pt, MAG4*, and *MAG4/Pt* airways. Scale bars for each genotype are 300 μm (left) and 75 μm (right).

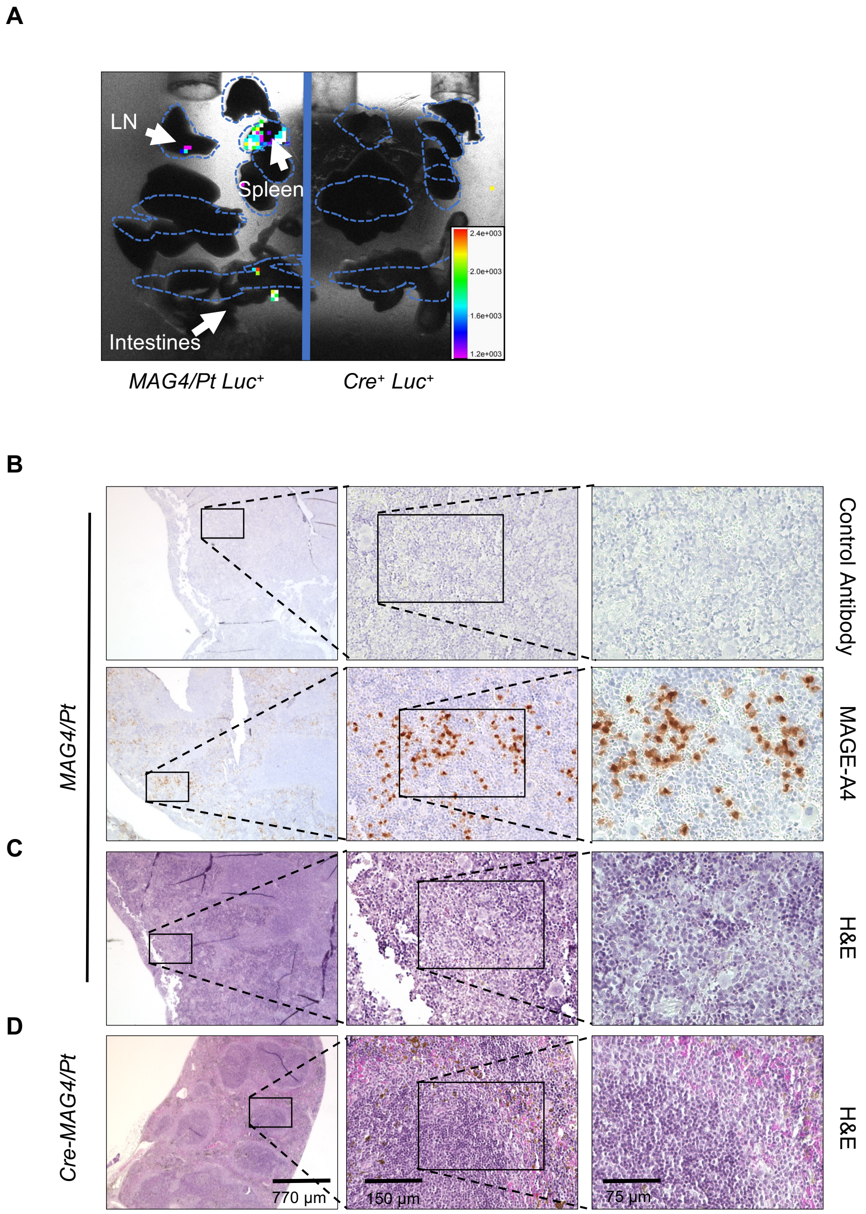

**Fig. S4: *MAG4/Pt* lung tumors metastasize**. (**A**) Representative luciferase detection i.e. of *MAG4/Pt Luc* at 13 months of age showing distal metastasis, LN: lymph node. Other organs are stomach, kidneys, and liver (4 independent experiments). (**B**) Secondary (control) antibody alone and MAGE-A4 IHC staining of *MAG4/Pt* spleen. (**C**) Respective H&E of *MAG4/Pt* spleen from (B). Rightmost images in (B-D) are also shown in Figure 2E. (**D**) H&E of *Cre- MAG4/Pt* spleen. (**B-D**) The scale bars from left to right are: 770 µm, 150 µm, 75 µm.

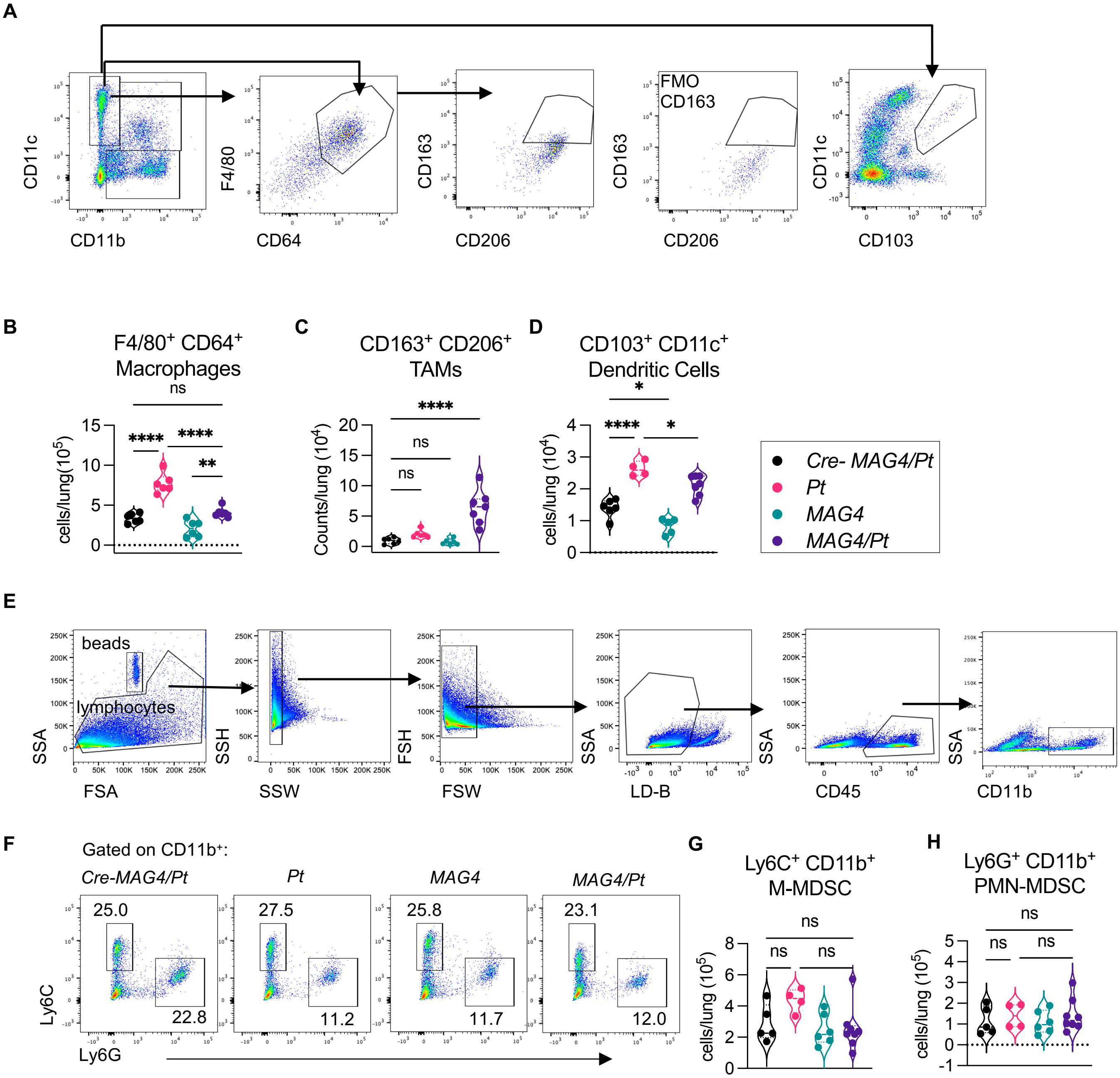

**Fig. S5: Innate immune cells flow cytometry gating strategies and representative plots show expansion of tumor-associated macrophages and inhibition of antigen-presenting cells in *MAG4/Pt* mice.** (**A**) Flow cytometric analyses from 5-month-old mouse whole-lung single cell suspension showing cell subsets gating strategy. Cell counts of (**B**) F4/80$^+$ CD64$^+$ CD11c$^+$ CD11b$^-$ macrophages, (**C**) CD163$^+$ CD206$^+$ Tumor associated macrophages (TAMs), and (**D**) CD103$^+$ CD11c$^+$ Dendritic Cells. n = 5-7 mice per group, repeated 2 times in different groups of mice; (**E**) Gating strategy for lung myeloid-derived suppressor cell (MDSC) populations in 8-month-old mouse whole lung (**F**) Representative images and quantification of (**G**) Ly6C$^+$ mononuclear (M)-MDSCs and (**H**) Ly6G$^+$ polymorphonuclear (PMN)-MDSCs. Representative of 2 independent experiments, n= 4-9 per group. One-way ANOVA Tukey's correction. *$P$<0.05, **$P$<0.01, ***$P$<0.001, ****$P$<0.0001

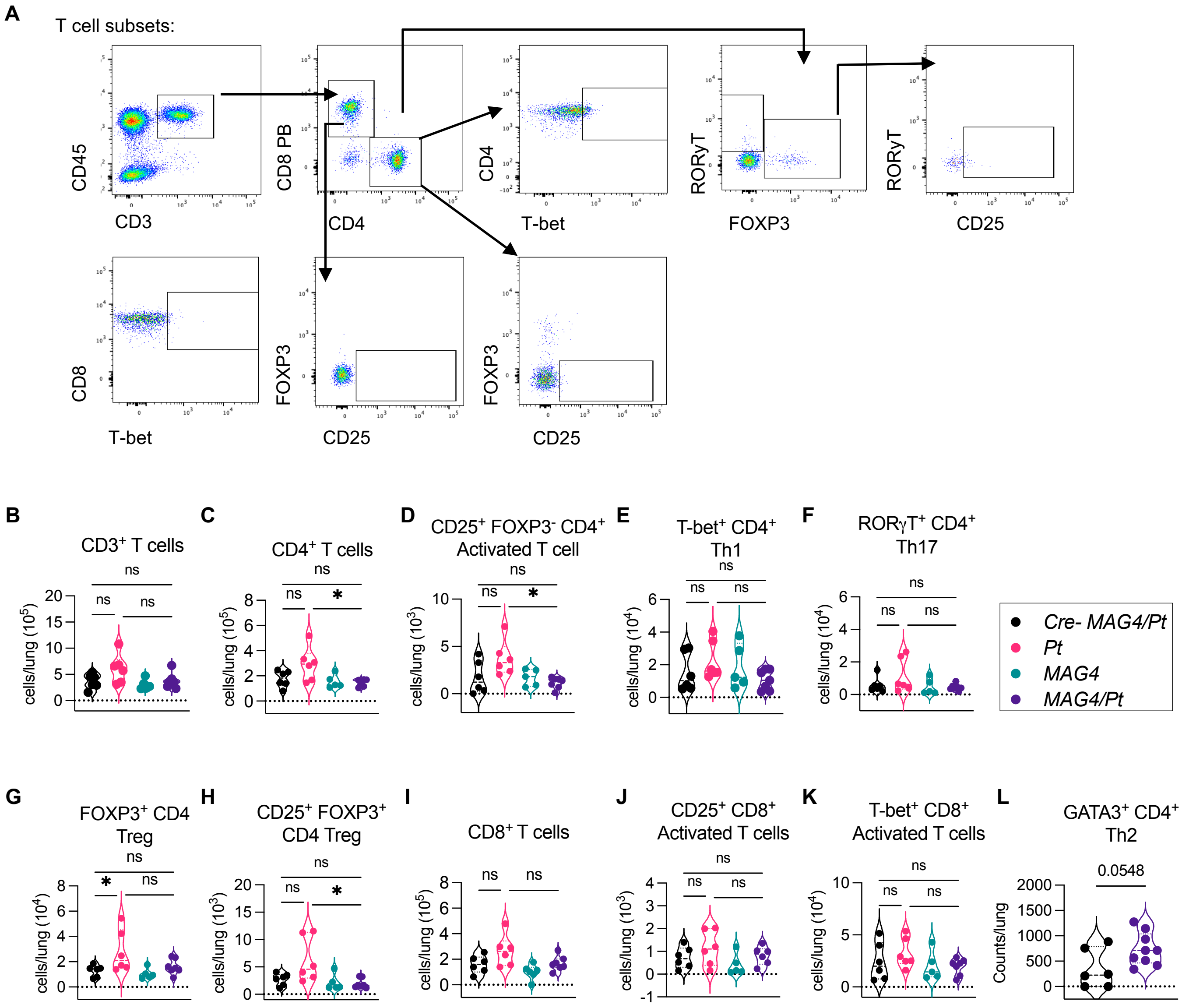

**Fig. S6: Reduced CD4 T cell subsets in MAG4/Pt mice.** (**A**) Flow cytometric analyses from 5-month-old mouse whole-lung single cell suspension gating strategy for T cell subsets (**B**-**K**). (**B**) Total $CD3^+$ T cells (**C**) Total $CD4^+$ T cells and CD4 subsets: (**D**) $CD25^+$ $FOXP3^-$ $CD4^+$ Activated T cells, (**E**) T-$bet^+$ (Th1) (**F**) $ROR\gamma T^+$ (Th17), (**G**) $FOXP3^+$ (Treg) and (**H**) $CD25^+$ on same Treg cells (from (**G**)), and. (**I**) Total $CD8^+$ T cells and subsets (**J**) $CD25^+$ and (**K**) T-$bet^+$ as indicators of activated $CD8^+$ T cells. n = 5-7 mice per group, one experiment. (**L**) Total Th2 ($GATA3^+$ CD4) T cells n = 6-9 per group, one experiment, mice age 7-8 months. One-way ANOVA Tukey's correction or student t-test. *$P$<0.05, ns = not significant

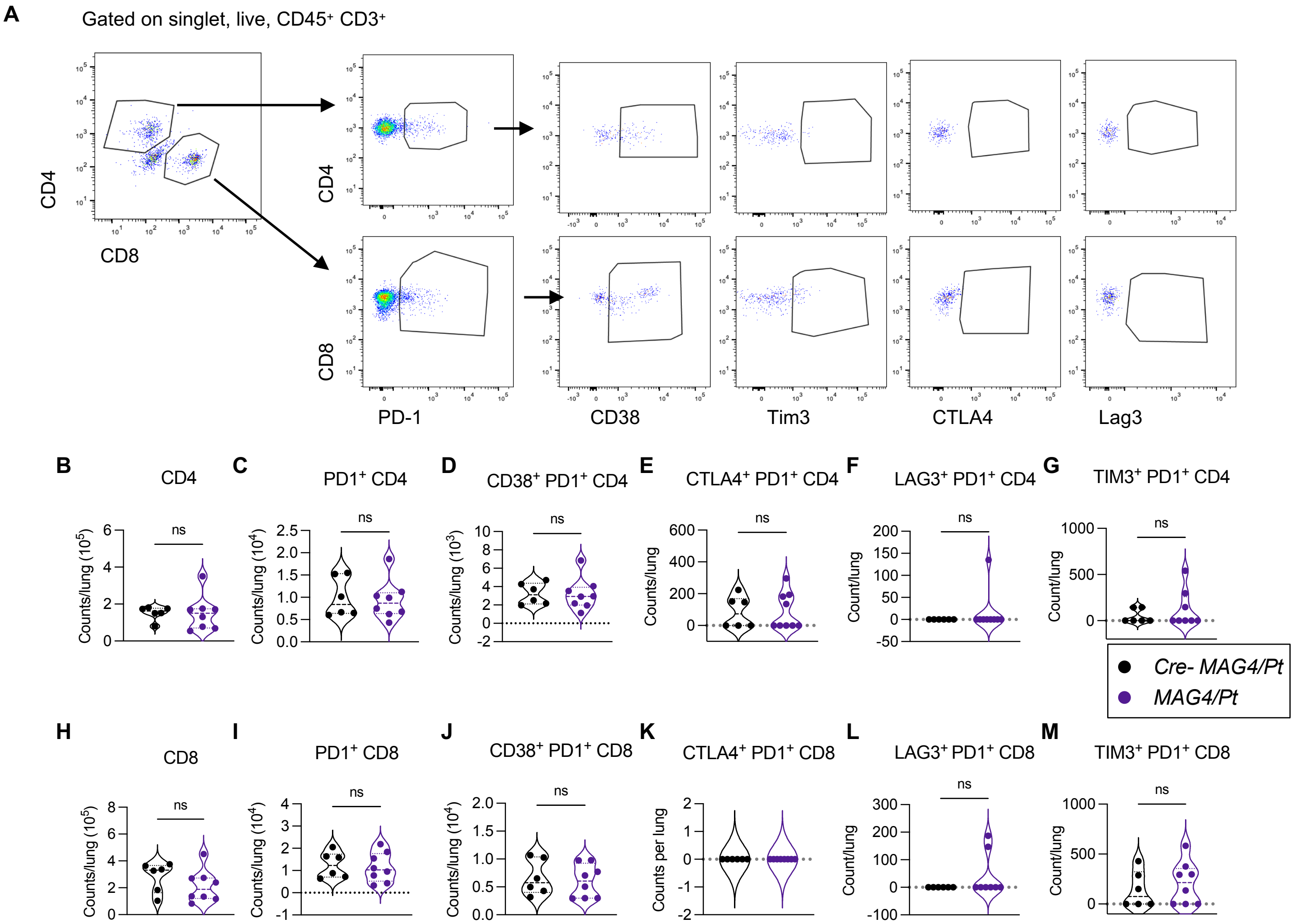

**Fig. S7: T cell exhaustion markers flow cytometry gating strategies and quantification.** (**A**) Flow cytometric analysis of lung from 7–8-month-old mice T cell gating strategy for immunosuppressive markers. Quantification of CD4 T cells and (**C**) PD-1$^+$ then co-expression of PD-1 with (**D**) CD38, (**E**) CTLA4, (**F**) LAG3, or (**G**) TIM3. (**H**) Quantification of CD8 T cells and (**I**) PD-1$^+$ then co-expression of PD-1$^+$ CD8 T cells with (**J**) CD38, (**K**) CTLA4, (**L**) LAG3, or (**M**) TIM3. Significance determined by student t-test. NS= not significant.

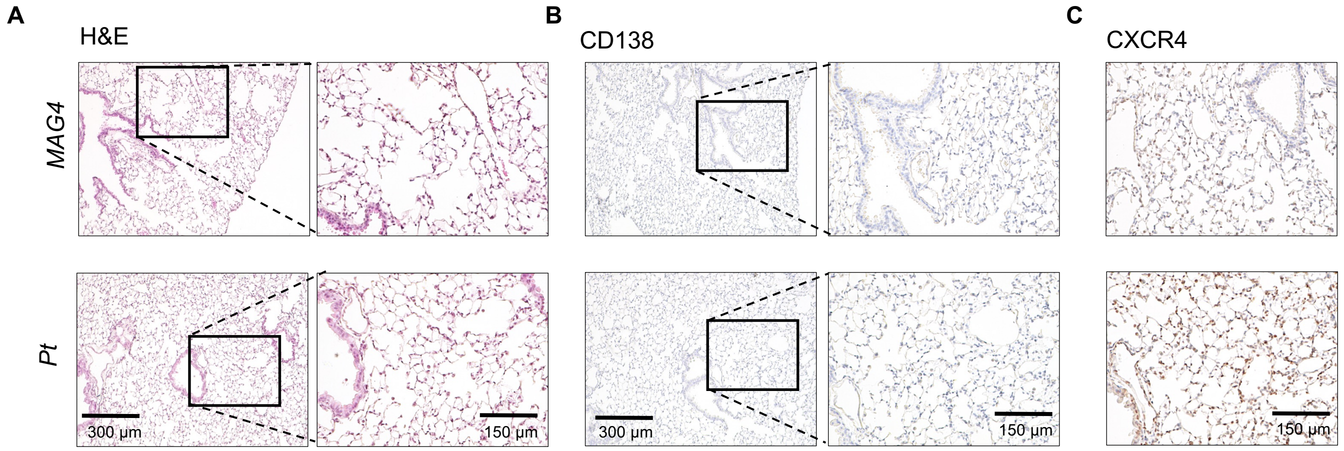

**Fig. S8: CD138 and CXCR4 staining of *MAG4* and *Pt* lung** (**A**) Representative serial section lung H&E from 5-month-old *MAG4* and *Pt* mouse. Scale bars represent 300 µm (left) and 150 µm (right) . (**B-C**) Representative serial sections from (A), IHC of (**B**) CD138; scale bars are 300 µm (left) and 150 µm (right), and (**C**) CXCR4 from 5-month-old *MAG4* and *Pt* mouse; the scale bar is 150 µm.

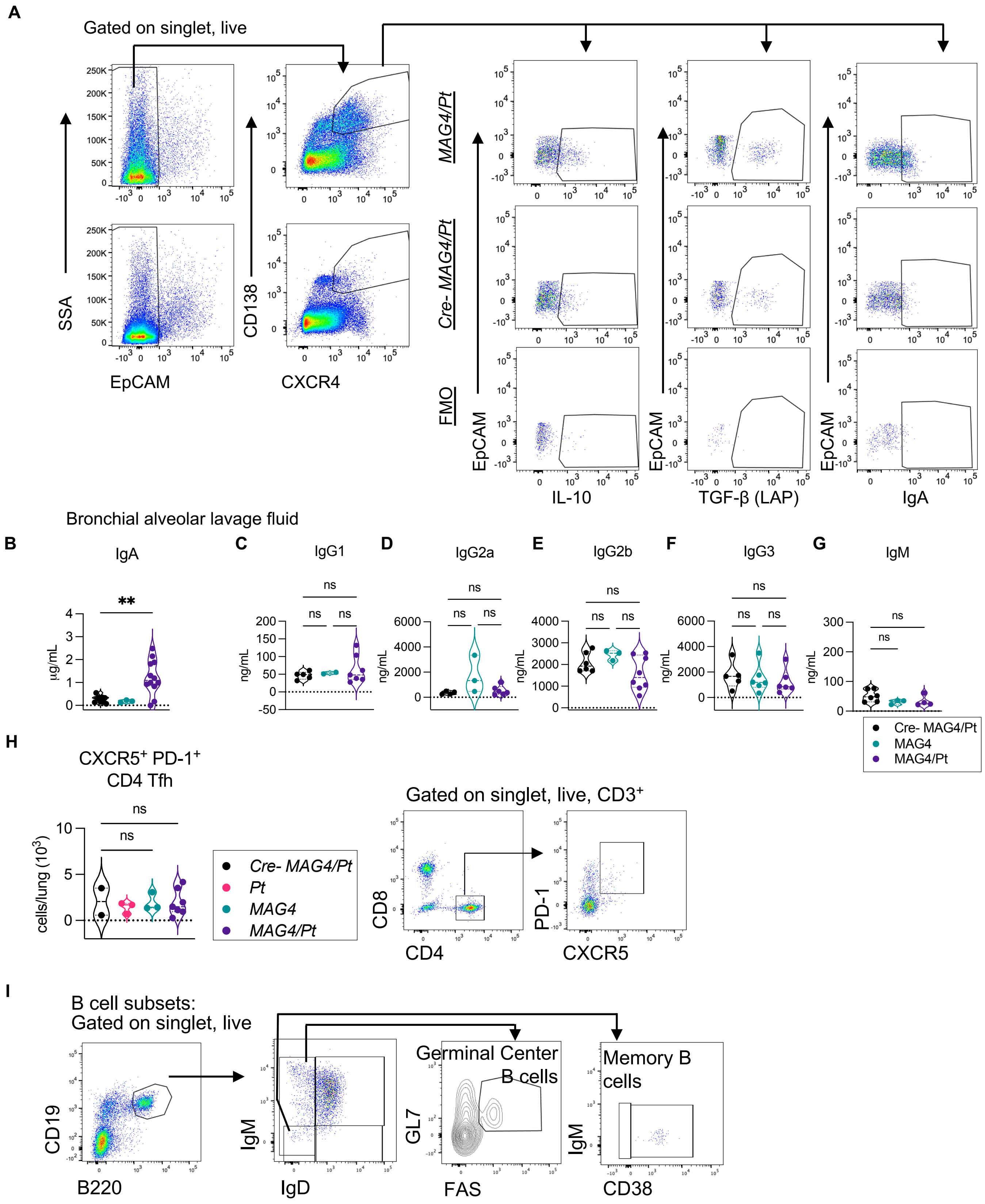

**Fig. S9: Plasma cell and B cell characteristics and gating strategies**. (**A**) Plasma cell gating strategy and representative gating for IgA, TGF-β (measured by latent activating peptide; LAP), and IL-10. FMO= fluorescence minus one. (**B-G**) Immunoglobulin isotype in bronchial alveolar lavage fluid (BAL) from 7-month-old mice as detected by ELISA, representative of 2-3 experiments, n=3-12 per group. (**B**) Immunoglobulin A (IgA), (**C**) Immunoglobulin G1 (IgG1), (**D**) Immunoglobulin G2a (IgG2a), (**E**) Immunoglobulin G2b (IgG2b), (**F**) Immunoglobulin G3 (IgG3), (**G**) Immunoglobulin M (IgM). (**H**) Tfh ($CXCR5^+$ $PD\text{-}1^+$ $CD4^+$ T cells) lung counts (left) measured by flow cytometry in 5-month-old mice and gating strategy (right). (**I**) Gating strategy for B cell subsets. Significance determined by one-way ANOVA with Tukey or Dunnett's correction, ***P* <0.01, ****P* <0.001, *****P* <0.0001, ns = not significant.

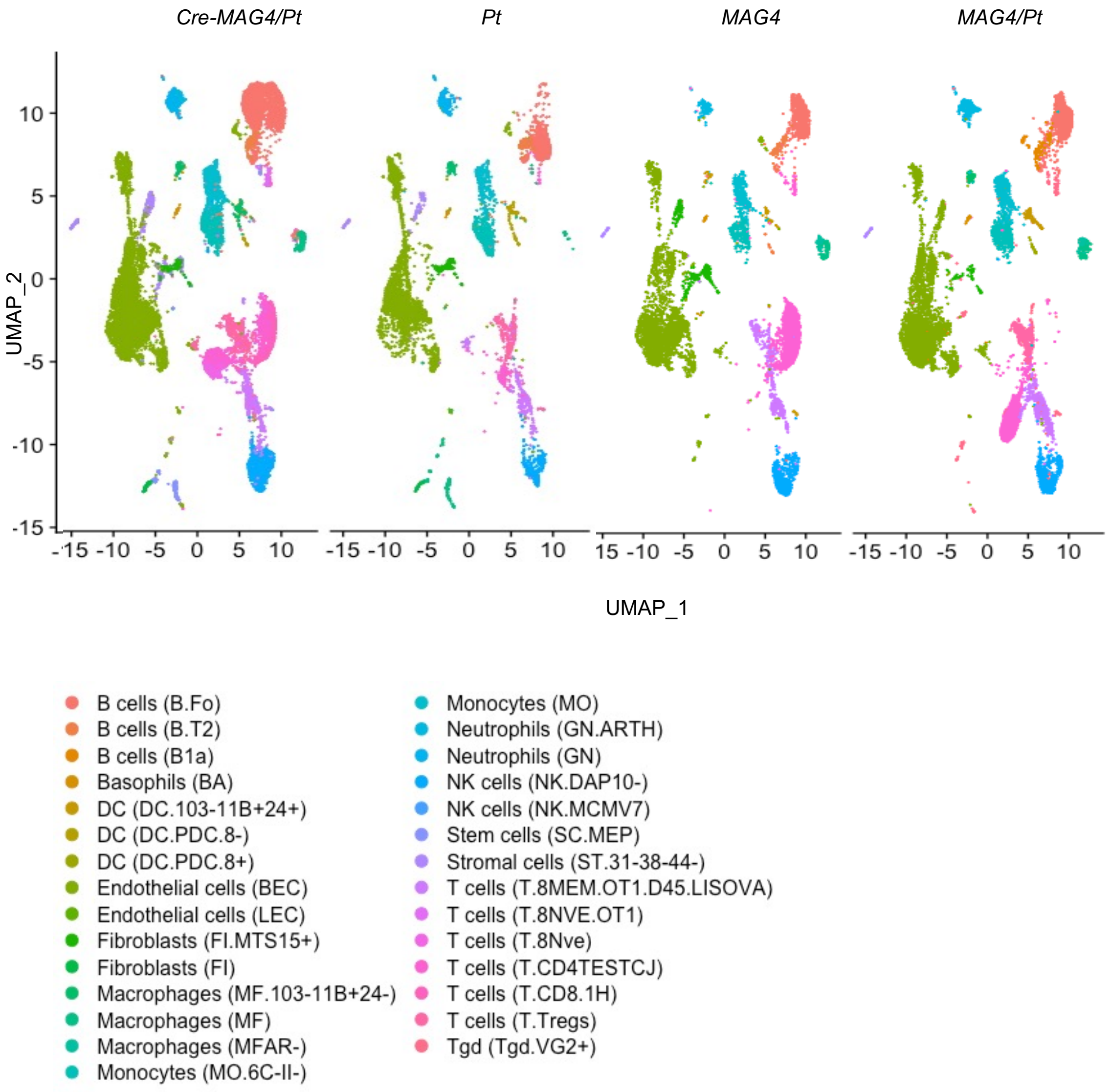

**Fig. S10: scRNA seq distribution of cell clusters by genotype.** Featureplots showing whole-lung of 5-month old mice clustered by cell type separated by individual genotype.

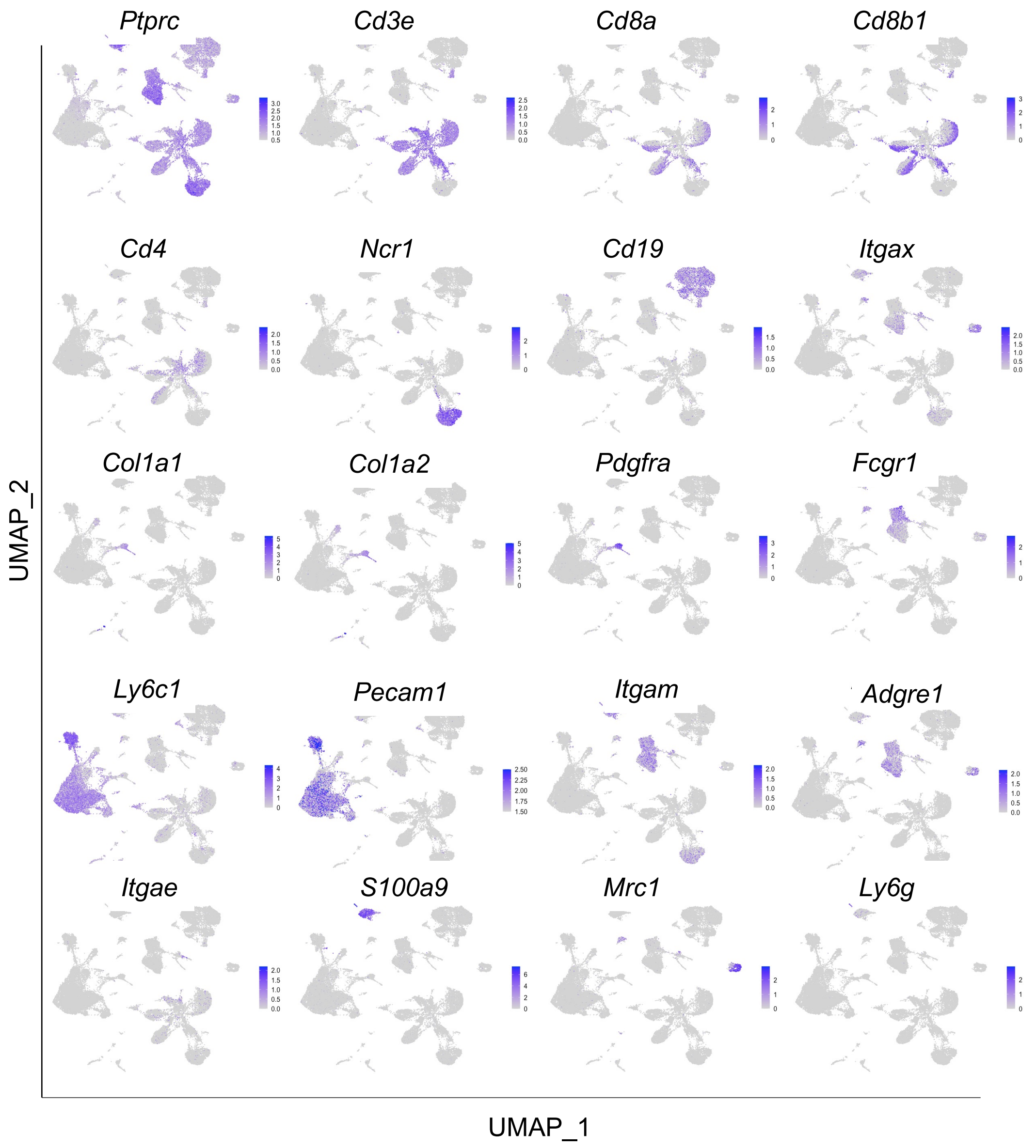

**Fig. S11: scRNA seq cluster identification**. Featureplots showing expression patterns of known cell identity markers to verify identity of clusters determined by SingleR.

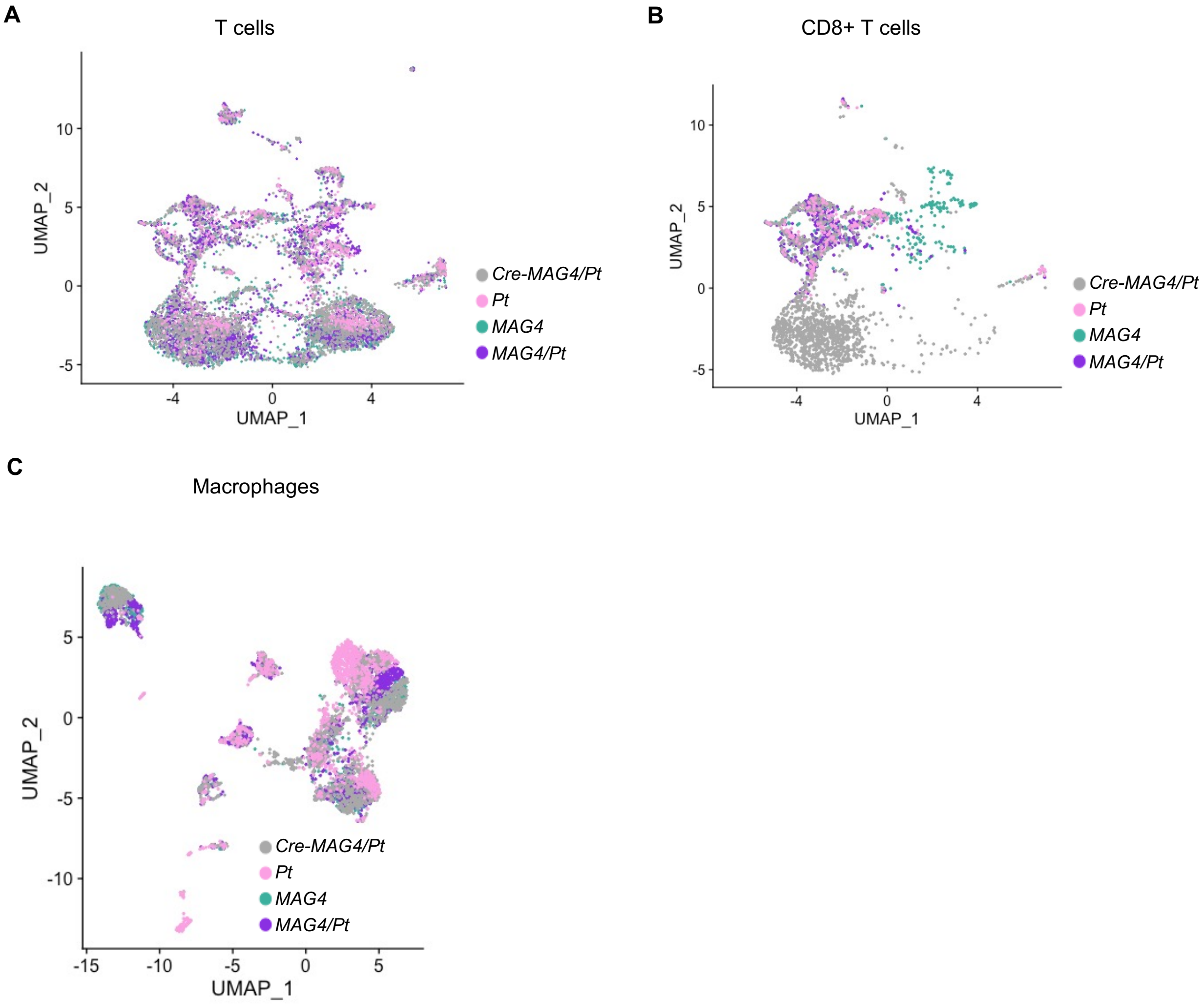

**Fig. S12: scRNA seq assessment of immune populations.** (**A**) UMAP showing distribution of all T cell populations by genotype. (**B**) UMAP showing distribution of CD8+ T cells by genotype. (**C**) UMAP projecting distribution of macrophages by genotype.

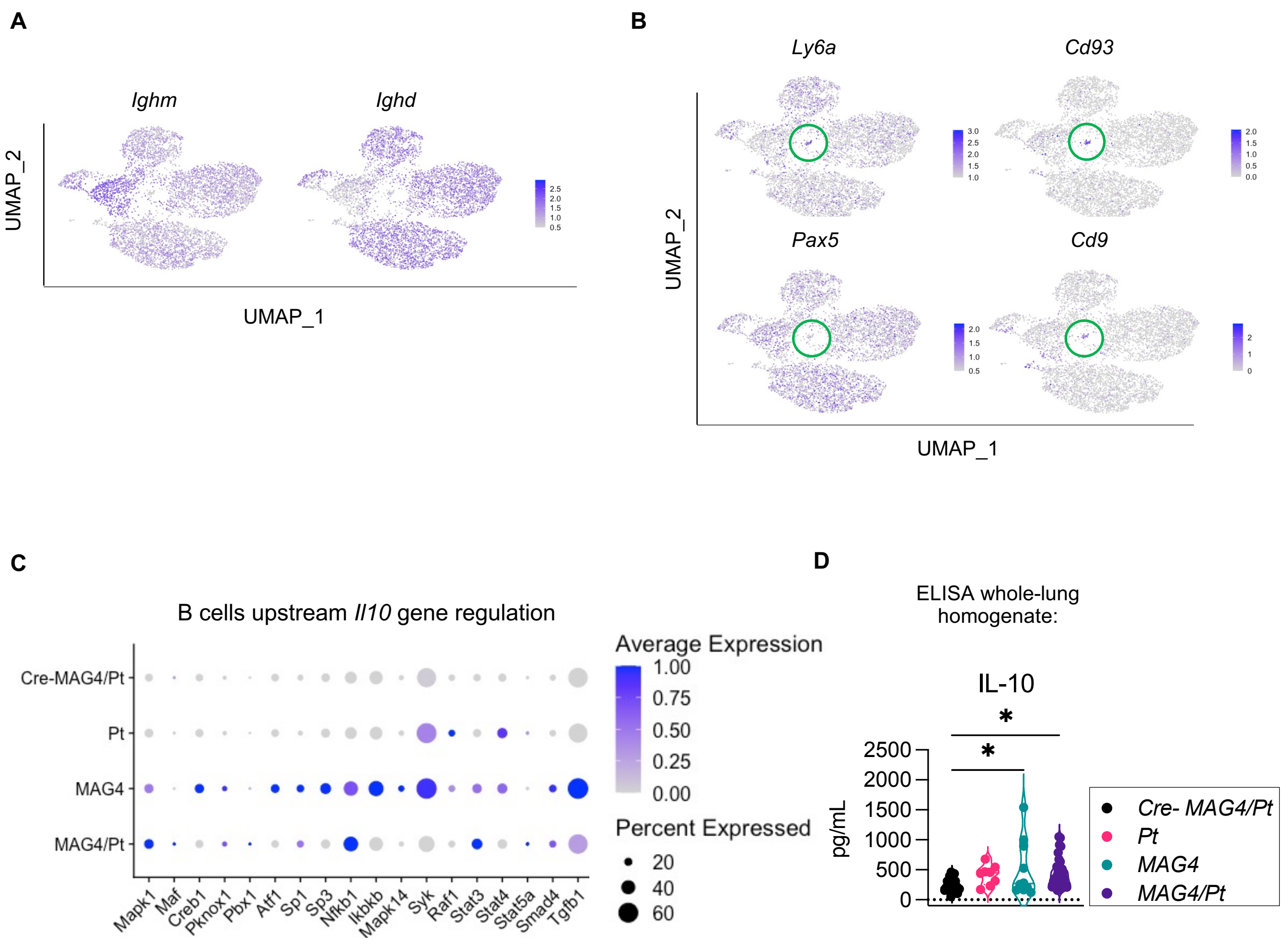

**Fig. S13: B cell characteristics** (**A**) scRNA-seq featureplots show by distribution that naïve B cells are double-positive for *Ighm* and *Ighd*. (**B**) scRNA-seq featureplots show center cluster increased expression of *Ly6a*, *Cd93*, and *Cd9* and lack of *Pax5.* (**C**) scRNA-seq dotplot comparing relative expression levels of genes known to induce *Il10* in various immune cell types and of *Tgfβ1* among the genetic conditions (**D**) ELISA of IL-10 of supernatant collected from overnight whole-lung homogenate culture, 2 experiments combined, n=8-32 per group. *$P$ < 0.05, One-way ANOVA with Dunnett's correction.

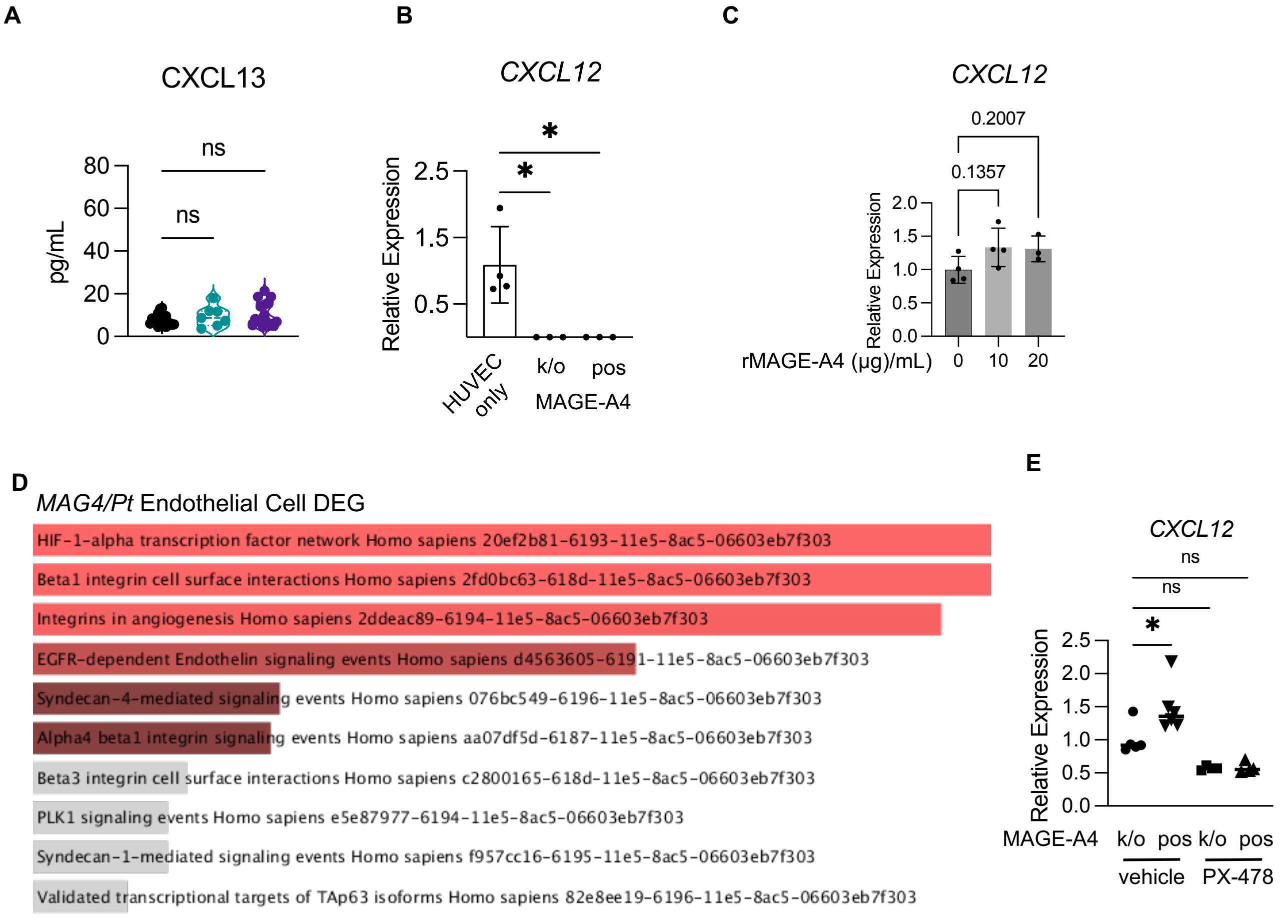

**Fig. S14. *CXCL12* specific to endothelial cells**. (**A**) CXCL13 concentration from whole-lung homogenate from 7-8 months old mice by ELISA. Data from 3 independent experiments combined, n=7-21 per group (**B**) Baseline difference in *CXCL12* by qPCR between human endothelial cells (HUVECs) and H1299 NSCLC cells cultured separately. (**C**) Relative expression of *CXCL12* in HUVECs when treated with recombinant (r)MAGE-A4 at the indicated concentrations for 24 hours. (**D**) Pathway analysis using NCI database from Enrichr online software of differentially expressed genes (DEG) from *MAG4/Pt* endothelial cells compared to those from *Cre-MAG4/Pt* (**E**) *CXCL12* relative expression in co-culture of HUVECs with MAGE-A4-positive (pos) or knock-out (k/o) H1299 cells treated with vehicle (DMSO) or PX-478 at 10μM for 24 hours. n=4-6 per group, representative of 2 independent experiments. Significance determined by One-way ANOVA with Dunnett's correction.
*$P$ < 0.05

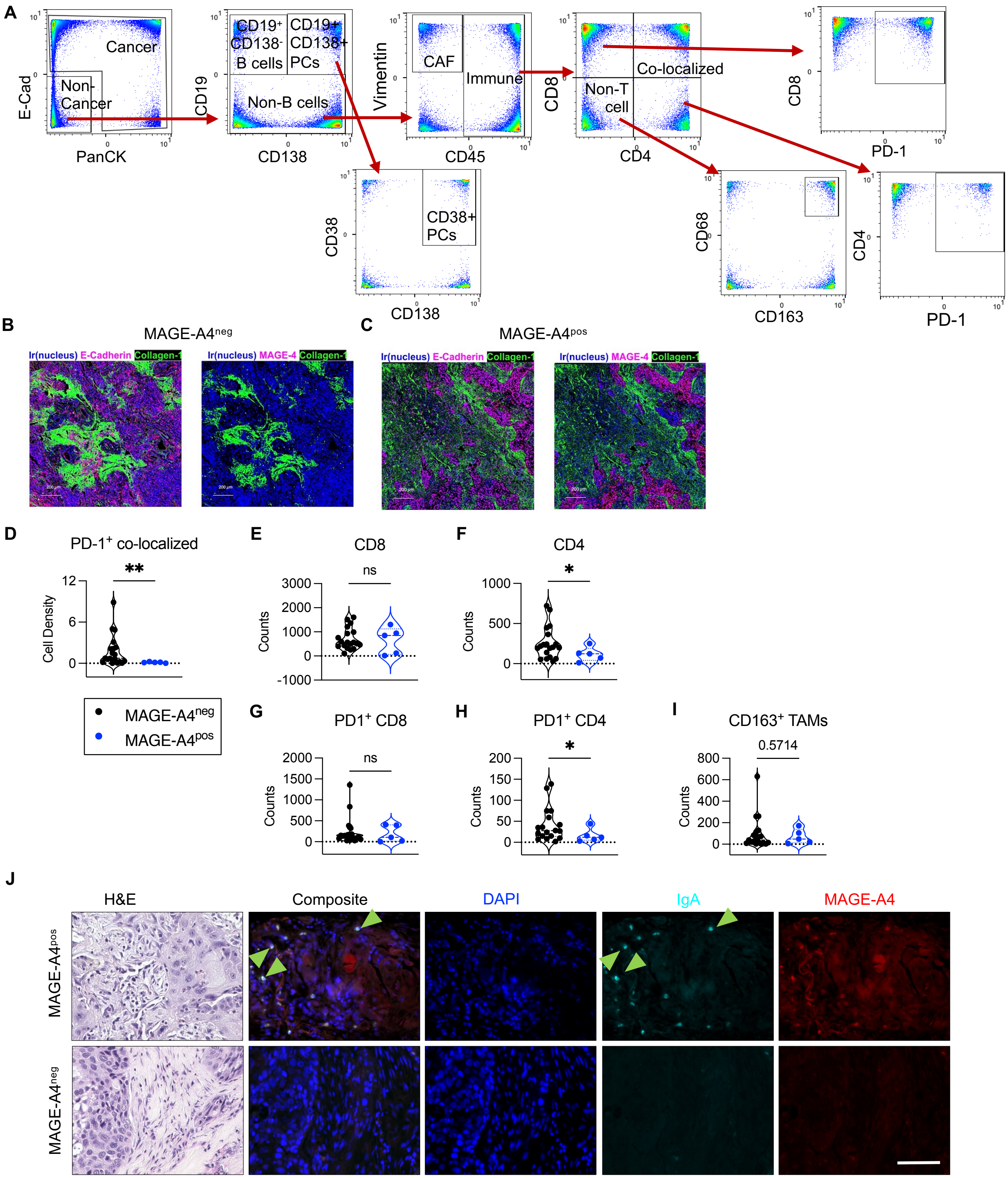

**Fig. S15: Imaging mass cytometry reveals T cells are abrogated in MAGE-A4pos tumors**. Imaging Mass Cytometry was utilized to evaluate a human tumor microarray of 24 individual NSCLC samples. (**A**) Gating strategy for phenotyping cells generated by IMC. PCs = plasma cells, CAF = Cancer Associated Fibroblast. (**B**, **C**) Representative image of MAGE-A4neg tumor (**B**) and MAGE-A4pos (**C**) showing co-localization of E-Cadherin (magenta, left image) with MAGE-A4 (magenta, right image). **(D)** Cell density quantification of PD-1+ CD8:CD4 co-localized T cells.(**E**) CD8 and (**F**) CD4 T cell counts in MAGE-A4neg and MAGE-A4pos NSCLC samples. (**G**) Quantification of PD-1+ CD8 and (**H**) PD-1+ CD4 T cells. (**I**) CD163+ CD68+ tumor associated macrophages (TAMs). (**J**) Representative immunofluorescent staining of IgA and MAGE-A4 in MAGE-A4pos (top) and MAGE-A4neg (bottom) lung squamous cell carcinoma. Green arrows indicate IgA-expressing cells. Scale bar = 75 µm. Significance determined by Welch's or student t-test. *$P < 0.05$, **$P < 0.01$, ns= not significant.

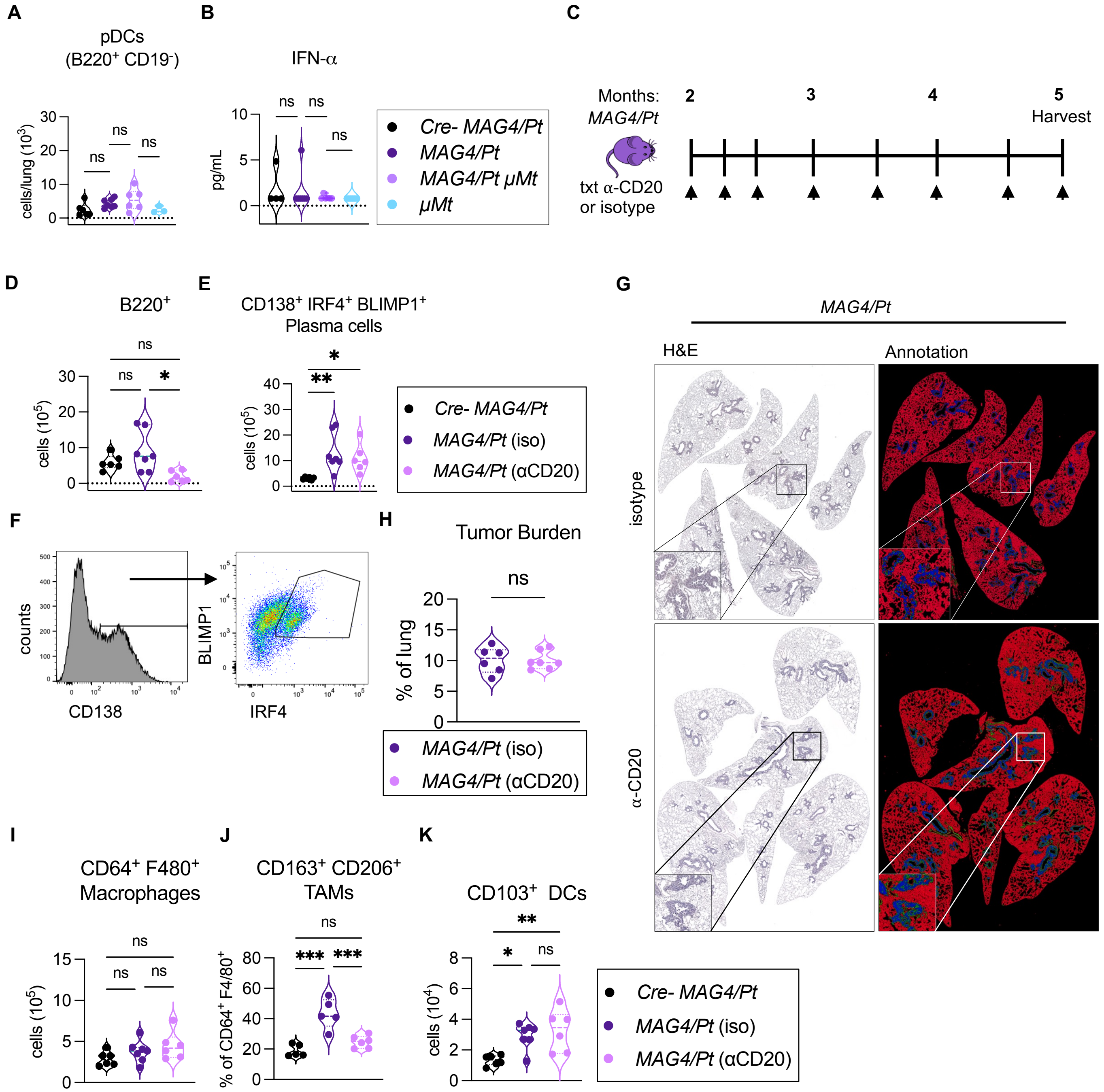

**Fig. S16: B cell depletion is not sufficient to reduce tumor burden in *MAG4/Pt*.** (**A**) Flow cytometric analysis whole lung for B220+ CD19- plasmacytoid dendritic cells (pDCs). (**B**) Quantification of IFN-α in bronchial alveolar lavage fluid by LEGENDPlex. *N*=3-7 per group (**C**) Schematic of α-CD20-mediated B cell depletion. Beginning at 2 months of age, *MAG4/Pt* mice were injected i.p. with α-CD20 antibody or isotype control once per week for 3 weeks then bi-weekly up to 5 months of age. (**D-H**) Flow cytometric analyses in right 3 lobes and post-caval lobe. Cell counts are reflected as total cells of these lobes. (**D**) Validation of effective B cell depletion of B220+ B cells as counts and (**E**) Plasma cell (CD138+ IRF4+ Blimp1+) quantification in treated and untreated lung. (**F**) Gating strategy for plasma cells in (**E**). (**G**) Representative H&E (left) and Avia software annotation (right) of *MAG4/Pt* lung treated with isotype or α-CD20. (**H**) Quantification of lung tumor burden as percentage of total lung area. n= 6-7 per group, one experiment. (**I**) Total CD64+ F4/80+ macrophages and (**J**) relative abundance of CD163+ CD206+ Tumor Associated Macrophages (TAMs) (**K**) Total CD103+ Dendritic cells. Significance was determined by One-way ANOVA with Tukey's correction or student t-test. *$P < 0.05$, **$P < 0.01$, ***$P < 0.001$, ****$P < 0.0001$, ns= not significant.

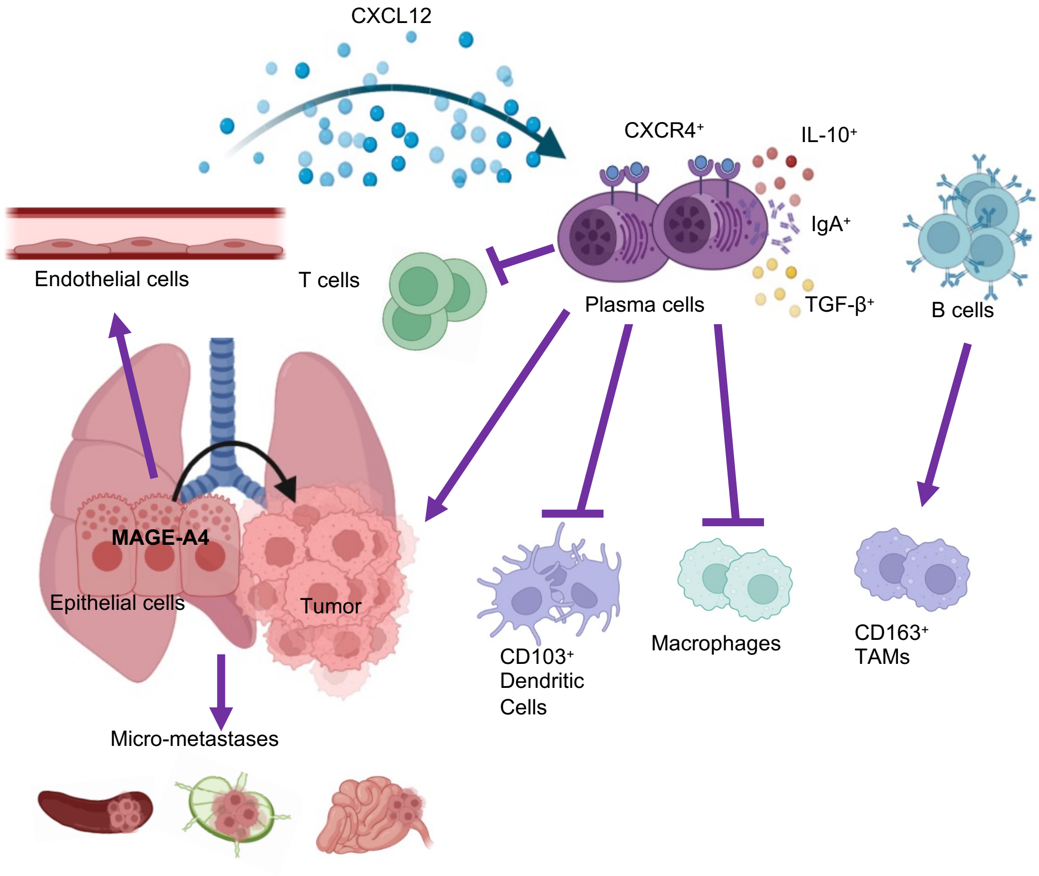

**Graphical Abstract: MAGE-A4 role in NSCLC development by immune modulation.** MAGE-A4 promotes tumorigenesis and metastasis. MAGE-A4 stimulates endothelial cells to increase production of CXCL12, which recruits and retains cognate $CXCR4^+$ plasma cells, which express TGFβ and IL-10 and produce IgA. These immunosuppressive plasma cells inhibit antigen presenting cells, including macrophages and cross-presenting $CD103^+$ Dendritic cells. Immunosuppressive plasma cells also inhibit the infiltration and activation of all T cell subsets. While $CD163^+$ tumor associated macrophages (TAMs) are increased in the *MAG4/Pt* tumor microenvironment, these are driven by B cells, and the absence of both immune cell types does not affect tumor burden. In contrast, plasma cells promote tumor progression.

Supplemental Table 1: NSCLC patient demographics and characteristics

| Lab ID | Tumor Type | Age | Gender | Race | Smoking Status | Pack/year |
|---|---|---|---|---|---|---|
| 01 | adenocarcinoma | 61 | M | C | Current | 100 |
| 02 | adenocarcinoma | 68 | M | AA | Current | 100 |
| 03 | adenocarcinoma | 79 | M | C | Former | 30 |
| 04 | adenocarcinoma | 70 | M | C | Current | 60 |
| 05 | adenocarcinoma | 72 | M | AA | Current | 54 |
| 05 | adenocarcinoma | 55 | M | AA | Former | 10 |
| 07 | adenocarcinoma | 74 | M | C | Current | 52 |
| 08 | adenocarcinoma | 73 | M | C | Former | 50 |
| 09 | adenocarcinoma | 74 | M | AA | Current | 50 |
| 10 | adenocarcinoma (mucin) | 73 | M | AA | Former | 52 |
| 11 | adenocarcinoma (mucin) | 77 | M | C | Current | 60 |
| 12 | Adenosquamous ca | 74 | M | C | Current | 50 |
| 13 | Adenosquamous ca | 65 | M | C | Current | 25 |
| 14 | Adenosquamous ca | 63 | F | AA | Current | 40 |
| 15 | large cell | 69 | M | C | Current | 50 |
| 16 | large cell (sarcomatoid pattern) | 57 | M | C | Current | 55 |
| 17 | squamous cell ca | 75 | M | AA | Current | 30 |
| 18 | squamous cell ca | 66 | M | C | Never | 0 |
| 19 | squamous cell ca | 76 | M | C | Former | 50 |
| 20 | squamous cell ca | 63 | M | C | Former | 100 |
| 21 | squamous cell ca | 68 | M | H | Former | 100 |
| 22 | Squamous cell ca | 69 | M | C | Current | 112 |
| 23 | squamous cell ca | 83 | M | AA | Former | 50 |
| 24 | squamous cell ca | 64 | M | C | Former | 50 |
| 25 | squamous cell ca | 73 | M | AA | Current | 50 |
| 26 | squamous cell ca | 61 | M | C | Current | 100 |
| 27 | squamous cell ca | 69 | M | C | Current | 55 |
| 28 | squamous cell ca | 69 | M | C | Current | 53 |
| 29 | squamous cell ca (basaloid) | 62 | M | C | Never | 0 |

Supplemental Table 2: Flow Cytometry Antibodies

| Target | Fluorophore | Vendor | Clone | Catalog # |
|---|---|---|---|---|
| CD3 | PerCP-Cy5.5 | Biolegend | 17A2 | 100218 |
| CD4 | FITC | Biolegend | RM4-5 | 100510 |
| CD8 | PB | Biolegend | 53-6.7 | 100725 |
| PD-1 | PE-Cy7 | Biolegend | RPM1-30 | 109110 |
| PD-1 | APC-R700 | BD Biosciences | J43 | 565815 |
| LAG3 | APC | Biolegend | C9B7W | 125210 |
| CTLA4 | BV605 | Biolegend | UC10-4B9 | 106323 |
| TIM3 | BV650 | BD Biosciences | 5D12/Tim3 | 747623 |
| CD38 | PE-Cy7 | Biolegend | 90 | 102718 |
| F4/80 | BUV395 | BD Biosciences | T45-2342 | 565614 |
| CD19 | APCR700 | BD Biosciences | ID3 | 565473 |
| LAP (TGF-β1) | PE | Biolegend | TW7-16B4 | 141404 |
| IL-10 | APC | Biolegend | JES5-16E3 | 505010 |
| CD206 | PE | Biolegend | C068C2 | 141706 |
| Gata3 | PE | Biolegend | 16E10A23 | 653804 |
| CD45 | BV510 | Biolegend | 30-F11 | 103138 |
| CD45.2 | AF488 | Biolegend | 104 | 109816 |
| B220 | AF488 | Biolegend | RA3-6B2 | 103225 |
| T-bet | BV650 | BD Biosciences | O4-46 | 564142 |
| IgA | BV650 | BD Biosciences | C10-1 | 743296 |
| IgA | FITC | Invitrogen | mA-6E1 | 11-4204-81 |
| CD11b | PE-Cy7 | Biolegend | M1/70 | 101216 |
| CD24 | PerCP-CY5.5 | Biolegend | M1/69 | 101823 |
| CD103 | APC | Biolegend | 2E7 | 121414 |
| CD163 | FITC | eBiosciences | TNKUPJ | 11-1631-82 |
| CD11c | PB | Biolegend | N418 | 117322 |
| IRF4 | PE | Biolegend | IRF4.3E4 | 646403 |
| CXCR4 | BV421 | Biolegend | L276F12 | 146511 |
| CD19 | BV510 | Biolegend | 6D5 | 115545 |
| CD138 | BV605 | Biolegend | 281-2 | 142531 |
| CD138 | PE | Biolegend | 281-2 | 142504 |
| Blimp1 | APC | Biolegend | 5E7 | 150007 |
| BCMA (CD269) | PE-Vio770 | Miltenyi | REA550 | 130-108-325 |
| IgM | BV510 | BD Biosciences | AF6-78 | 742344 |
| GL7 | PB | Biolegend | GL7 | 144614 |
| CD95 (FAS) | PE | Biolegend | SA367H8 | 152608 |
| IgD | AF647 | Biolegend | 11-26c.2a | 405708 |
| CD38 | PE-Cy7 | Biolegend | 90 | 162718 |
| CD25 | BB515 | BD Biosciences | PC61 | 564424 |
| FOXP3 | APC | Invitrogen | FJK-16s | 17-5773-82 |
| RORγT | PE | BD Biosciences | Q31-378 | 562607 |
| EpCAM | APC | Biolegend | G8.8 | 118214 |
| CXCR5 | BV605 | Biolegend | L138D7 | 145513 |

Supplemental Table 3: Imaging Mass Cytometry Antibodies and Conjugation

| Marker | Clone | Metal | Reactivity | BCM Lot | Dilutions (1:xx) |
|---|---|---|---|---|---|
| Vimentin | D21H3 | 110Cd | Mu/Hu/Rt/Mk | 210820B | 100 |
| Pan-CytoKeratin (C11) | C11 | 111Cd | Mu/Hu/Rt/Fi | 210820C | 50 |
| CD68 | KP1 | 112Cd | Hu | 210820D | 100 |
| Alpha-SMA | 1A4 | 141Pr | Hu/Mu/Rt | 201004A | 400 |
| CD19 | 6OMP31 | 142Nd | Hu/Mu/Rt | 210526A | 100 |
| CD223/LAG3 | D2G4O | 143Nd | Hu | 230306A | 50 |
| CD14 | EPR3653 | 144Nd | Hu/Mu | 210602D | 50 |
| T-bet | D6N8B | 145Nd | Hu | 210929E | 50 |
| CD16 | EPR16784 | 146Nd | Hu/Rt | 210601C | 50 |
| CD163/M130 | EDHu-1 | 147Sm | Hu | 221006A | 100 |
| MAGE-4 | E7O1U | 148Nd | Hu | 230222F | 50 |
| CD11b | EPR1344 | 149Sm | Hu/Mu/Rt | 210601E | 50 |
| PD-L1/CD274 | SP142 | 150Nd | Hu | 210811H | 100 |
| CD31 | EPR3094 | 151Eu | Hu | 210803D | 100 |
| CD45 | D9M8I | 152Sm | Hu | 210921H | 100 |
| CD44 | IM7 | 153Eu | Hu/Mu | 210826A | 200 |
| CD11c | EP1347Y | 154Sm | Hu/Cross | 210601D | 100 |
| FoxP3 | 236A/E7 | 155Gd | Hu | 210526B | 50 |
| CD4 | EPR6855 | 156Gd | Hu | 210811F | 50 |
| E-cadherin | 24E10 | 158Gd | Hu/Mu | 210929A | 50 |
| CD152/CTLA-4 | Ipilimumab | 159Tb | Hu | 230306C | 50 |
| CD38 | EPR4106 | 160Gd | Hu | 210617B | 100 |
| CXCL12 | D8G6H | 161Dy | Hu | 230222H | 50 |
| CD8a | C8/144B | 162Dy | Hu | 210526F | 50 |
| CD56 | EPR2566 | 163Dy | Hu | 210617D | 50 |
| CD15 | W6D3 | 164Dy | Hu | 210803C | 50 |
| PD-1/CD279 | EPR4877(2) | 165Ho | Hu | 210601B | 50 |
| CD45RA | HI100 | 166Er | Hu | 210526D | 50 |
| Granzyme B | EPR20129-217 | 167Er | Hu | 210819F | 50 |
| Ki-67 | 8D5 | 168Er | Hu/Mu | 220815B | 100 |
| Collagen 1 | 3G3 | 169Tm | Hu/Mu | 210601A | 100 |
| CD3 | D7A6E | 170Er | Hu | 210819C | 50 |
| CD138 | MI15 | 171Yb | Hu | 230222G | 100 |
| Cleaved Caspase-3 | 5A1E | 172Yb | Hu/Mu/Rt | 220509A | 50 |
| CD45RO | UCHL1 | 173Yb | Hu | 210526E | 200 |
| HLA-DR | YE2/36 HLK | 174Yb | Hu | 211116A | 100 |
| CD86 | BU63 | 175Lu | Hu | 210602C | 50 |
| CD33 | OTI2C1 | 176Yb | Hu | 210617E | 50 |
| Iridium | na | na | na | na | 1000 |

Supplemental Table 4: Immunoassay Antibodies

| **Western blot:** | | | | |
|---|---|---|---|---|
| **Target** | **Vendor** | **Dilution** | **Cat. #** | **Clone** |
| FLAG | Sigma-Aldrich | 1:3,000 | F7425 | SIG1-25 |
| Myc | Cell Signaling Technology | 1:1,000 | 2278 | 71D10 |
| MAGE-A4 | Santa-Cruz Biotechnology | 1:1,000 | sc-294929 | polyclonal |
| MAGE-A4 | Abcam | 1:1,000 | ab76177 | polyclonal |
| β-actin-HRP | Cell Signaling Technology | 1:5,000 | 12620S | D6A8 |
| Anti-rabbit HRP | Cell Signaling Technology | 1:10,000 | 7074S | |
| **IHC/IF** | | | | |
| **Target** | **Vendor** | **Dilution** | **Cat. #** | **Clone** |
| MAGE-A4 | Sigma | 1:1,000 (IHC) 1:50 (IF) | HPA021942 | |
| TTF1 | Santa Cruz Biotechnology | 1:200 | sc13040 | H-190 |
| KRT7 | Abcam | 1:8,000 | ab181598 | EPR17078 |
| Ki67 | Invitrogen | 1:200 | 14-5698-82 | SOLA15 |
| SOX2 | Milipore | 1:100 | ab5603 | |
| CD138 | ThermoFisher | 1:100 | 36-2900 | |
| KRT5 | Abcam | 1:200 | ab52635 | EP1601Y |
| CXCR4 | R&D | 8ug/mL | MAB21651 | 247506 |
| Biotinylated anti-rat | Vector Laboratories | 1:100 | BA-9400 | |
| Biotinylated anti-rabbit | Vector Laboratories | 1:400 | BA-5000 | |
| IgA | ThermoFisher | 1:100 | 3493-MSM1-P1 | HISA43 |
| Anti-mouse AF647 | ThermoFisher | 1:2,000 | A-21235 | |
| Anti-rabbit AF555 | ThermoFisher | 1:2,000 | A-21428 | |
| **ELISA Antibodies:** | | | | |
| Isotype (standard) | Vendor | Cat. # | | |
| IgG1 | Thermo Fisher Scientific | MG100 | | |
| IgG2a | Thermo Fisher Scientific | SA1-12079 | | |
| IgG2b | Thermo Fisher Scientific | MG2B00 | | |
| IgG3 | Thermo Fisher Scientific | MA7-10433 | | |
| IgM | Thermo Fisher Scientific | MGM00 | | |
| IgA | Southern Biotech | 016-01 | | |
